# Long-read RNA sequencing reveals allele-specific N^6^-methyladenosine modifications

**DOI:** 10.1101/2024.07.08.602538

**Authors:** Dayea Park, Can Cenik

**Affiliations:** Department of Molecular Biosciences, University of Texas at Austin, Austin, TX 78712, USA

**Keywords:** LRS Special Issue, N^6^-Methyladenosine, Allele-specific expression

## Abstract

Long-read sequencing technology enables highly accurate detection of allele-specific RNA expression, providing insights into the effects of genetic variation on splicing and RNA abundance. Furthermore, the ability to directly sequence RNA promises the detection of RNA modifications in tandem with ascertaining the allelic origin of each molecule. Here, we leverage these advantages to determine allele-biased patterns of N^6^-methyladenosine (m6A) modifications in native mRNA. We utilized human and mouse cells with known genetic variants to assign allelic origin of each mRNA molecule combined with a supervised machine learning model to detect read-level m6A modification ratios. Our analyses revealed the importance of sequences adjacent to the DRACH-motif in determining m6A deposition, in addition to allelic differences that directly alter the motif. Moreover, we discovered allele-specific m6A modification (ASM) events with no genetic variants in close proximity to the differentially modified nucleotide, demonstrating the unique advantage of using long reads and surpassing the capabilities of antibody-based short-read approaches. This technological advancement promises to advance our understanding of the role of genetics in determining mRNA modifications.

## INTRODUCTION

Allele-specific expression (ASE) refers to the differences in gene expression from two alleles of the same gene. Such an imbalance in expression can contribute to phenotypic variation and the pathophysiology of diseases (Castel et al. 2015, 2020; Fan et al. 2020; de la Chapelle 2009; Gicquel et al. 2005). In mammalian development, a predominant form of ASE, genomic imprinting, plays a critical role as only one allele is expressed. Allele-specific DNA methylation and chromatin composition are two well-established epigenetic systems that control imprinted gene expression (Singh et al. 2010; Prendergast et al. 2012; Fournier et al. 2002). Particularly in development, DNA methylation regulates allele-specific expression, and coordinates X-chromosome inactivation in females for dosage compensation (Morcos et al. 2011; Benton et al. 2019). Furthermore, H3K27me3 marks in the early embryo mediate imprinted mono-allelic expression and persist from oocyte development through the blastocyst stage (Santini et al. 2021; Inoue 2023; Sergeeva et al. 2023).

ASE can reflect differential rates of transcription, mRNA stability, or alternative splicing between the alleles due to genetic variation (Amoah et al. 2021; Nembaware et al. 2008; Pai et al. 2012). That is, local genetic variants can influence transcriptional or post-transcriptional processes to modulate mRNA abundance of each allele (Robles-Espinoza et al. 2021). Intriguingly, while the significance of allele-specific RNA expression is well-acknowledged, allele-specific RNA modification remains underexplored.

N⁶-Methyladenosine (m6A), the most prevalent RNA modification of mRNAs, has been suggested to impact diverse mechanisms to regulate gene expression (He and He 2021; Lee et al. 2020). Various interactions with the methyltransferase complex or m6A reader proteins impact several steps of mRNA metabolism, including splicing, export, translation, recruitment of RNA binding proteins, and stability (Wang et al. 2022; Jiang et al. 2021; Lin and Gregory 2014; Akhtar et al. 2021).

Transcriptome-wide patterns of m6A RNA modifications have typically been studied using short-read sequencing coupled with either antibody-dependent methods such as MeRIP-seq (Meyer et al. 2012) or enzymatic/chemical approaches (Garcia-Campos et al. 2019; Meyer 2019a; Song et al. 2021). Among these methods, MeRIP-seq remains the most popular choice despite its limitations leading to elevated false-positive rates, attributable to nonspecific antibody binding (Helm et al. 2019; McIntyre et al. 2020; Zhang et al. 2021). Furthermore, all short-read sequencing strategies to detect m6A are inherently limited to aggregate measurements and are incapable of quantification at an individual molecular level.

In contrast, Oxford Nanopore Technology (ONT) RNA sequencing enables direct detection of RNA modifications such as m6A with single molecule resolution (Garalde et al. 2018). The electric signal recorded by the ONT sequencing platform was shown to be altered by the presence of RNA modifications (Workman et al. 2018; Garalde et al. 2018; Pelizzola et al. 2021). Subsequently, machine learning methods can utilize the electronic current signal intensity to identify potential m6A sites from long-read data (Hendra et al. 2022).

The ability to directly detect m6A modifications on the ONT RNA sequencing platform provides a unique opportunity to combine these advantages with the ability of long-read sequencing to facilitate ASE analysis. Long-read sequencing improves upon the fundamental limitations of short-read sequencing for allele-specific analysis by detecting an increased number of single nucleotide polymorphisms (SNPs) on a read, enabling its precise allelic assignment (Cho et al. 2014). This feature has been leveraged to characterize the genetic effects of rare and common variants in the transcriptome (Glinos et al. 2022). Furthermore, long-read sequencing enables comprehensive analysis of splicing (Tilgner et al. 2015, 2018; Joglekar et al. 2021) which has fundamental importance for determining mRNA modifications due to their dependence on splicing patterns and transcript architecture (Yang et al. 2022; He et al. 2023; Cenik et al. 2017).

Here we introduce a novel approach harnessing ONT direct RNA sequencing to surmount the persistent constraints of m6A detection methods for allele-specific analyses. Our findings establish long-read sequencing of RNA as a robust solution for allele-specific m6A modification (ASM) analysis.

## RESULTS

### ONT direct RNA sequencing enables the identification of allele-specific m6A modifications in hybrid mESCs

We leveraged ONT direct RNA sequencing to simultaneously determine the allelic origin of each molecule along with its m6A modification status. To achieve high accuracy of allelic assignment of individual molecules, we decided to use mouse embryonic stem cells (mESCs) that were derived from a cross of two highly genetically diverse mouse strains (C57BL/6J x CAST/EiJ, B6 x CAST) (Balasooriya and Spector 2022). Using ONT direct RNA sequencing, we generated two replicates of 2.3 and 2.2 million reads from these hybrid mESCs.

To assess our ability to accurately detect m6A modifications, we generated mESC clones where methyltransferase-like 3 (*Mettl3*), the major methyltransferase for m6A modifications, is knocked out (Bokar et al. 1994; Liu et al. 2013) (Supplemental Fig. 1A, Supplemental Table 1; Methods). As expected, in wild-type mESCs ∼6% of the adenines within the context of a DRACH motif had a high-probability (>0.85) of being modified, compared to only 0.2% in those with *Mettl3* knockout (Supplemental Fig. 1B). Among the sites displaying a high-probability modification ratio, the levels of modification ratios were consistently higher in wild-type compared to *Mettl3* knockout cells (Median modification ratio 0.629 and 0.512, respectively). Moreover, the modified adenines were predominantly clustered near the 3’ end of coding sequences, which is consistent with the expected pattern of m6A RNA modifications (Meyer et al. 2012; Zhang et al. 2019) (Fig. 1A-B; Supplemental Fig. 1C-D).

**Figure. 1.**
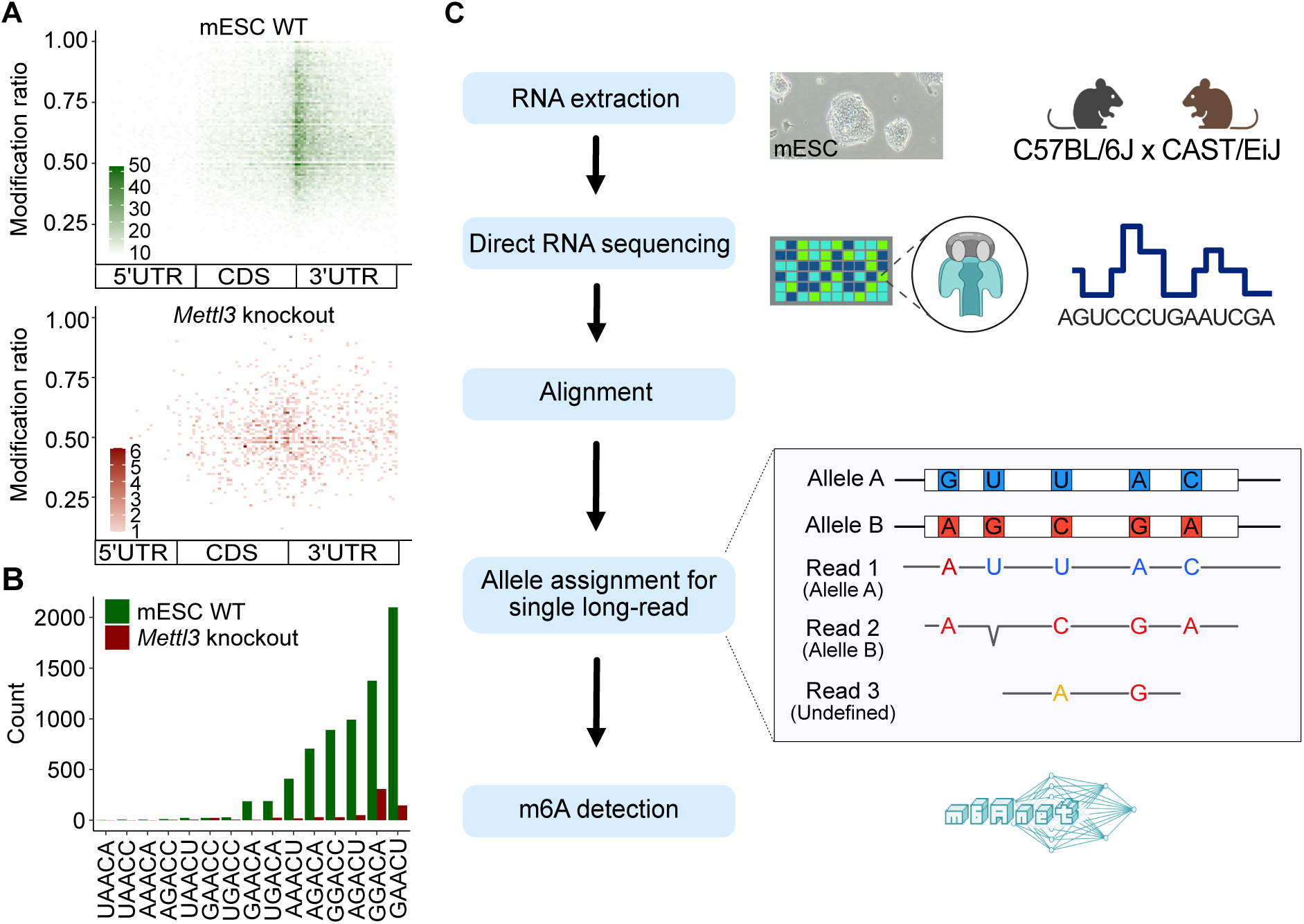
Allelic read assignment and m6A modification analysis using ONT direct RNA sequencing in hybrid mESCs. A) M6A modification ratio and locations detected from m6Anet using all reads (top, green, WT; bottom, red, *Mettl3* knockout). The relative m6A locations within the transcript body were determined. It presents modification ratios after high probability selection (> 0.85). The color darkness represents the counts of the ratio on the position. B) Comparison of the frequencies of instances of DRACH motif sequences (green, WT; red, *Mettl3* knockout). C) Schematic overview of the strategy used for allelic long-read assignment for allele-specific m6A modification analysis. Total RNA from hybrid mESC (C57BL/6J x CAST/EiJ) underwent ONT direct RNA sequencing. To avoid reference bias, we used an N-masked transcriptome for alignment. Reads were then allocated to each allele. This process classified reads into Allele A (B6), Allele B (CAST), and undefined categories, enabling m6A detection within each group individually.

Using single nucleotide polymorphisms (SNPs), we assigned 1,110,260 (replicate 1) and 837,011 (replicate 2) long-reads to their allelic origins across more than 13,000 transcripts (Fig. 1C, Methods). Of the detected transcripts, 8,657 transcripts had at least ten reads in both replicates. In allele-specific analyses, a common challenge is reference allele bias which is the tendency for reads that match the reference genome to align with a higher probability than reads containing the alternate allele, potentially skewing variant detection and analysis. To minimize this bias (Castel et al. 2015), we employed an N-masked transcriptome reference. This approach led to a mean CAST allele ratio across all transcripts of 0.505 as opposed to 0.485 when using an unmasked reference (Methods). These assignments were based on 135,380 (replicate 1) and 134,585 (replicate 2) informative positions on the long-reads that overlapped known genetic variation between the strains (210,004 total SNPs).

As an orthogonal approach to determine ASE, we used Illumina short-read sequencing. We found that RNA expression levels from the two methods were significantly correlated (Supplemental Fig. 2A; Spearman Correlations, 0.792-0.816 across replicates). Moreover, gene-level allele-specific RNA expression was moderately concordant between the two approaches (Supplemental Fig. 2B; weighted Spearman correlation coefficient 0.61, Methods). Although short-read sequencing produced nearly eight times more aligned reads, long-read sequencing identified 2.3 times as many SNPs, demonstrating that the greater number of informative positions in long-read data enhances allelic assignment accuracy and gene-specific ASE reproducibility (Spearman correlation coefficient 0.63 and 0.51 for long-read and short-read sequencing, respectively; Supplemental Fig. 2C-D). Taken together, these measures of quality control underscore the high precision in allelic assignment achieved through our methodology.

Following allelic read assignment, we employed a supervised machine learning approach (Hendra et al. 2022; Liu et al. 2019) to quantify m6A RNA modifications for the reads attributed to each allele. This process revealed an equivalence in the number of reads and m6A sites among alleles, indicating the allelic impartiality of our approach. Specifically, we observed similar numbers of reads (564,944 and 444,514 for B6; 557,787 and 443,131 for CAST) and potential modification sites (114,457 and 105,190 for B6; 112,947 and 105,117 for CAST) for each allele (Fig 2A, Supplemental Table 2). This result indicates minimal to no allelic bias in the assignment of reads and identification of modification sites.

**Figure. 2.**
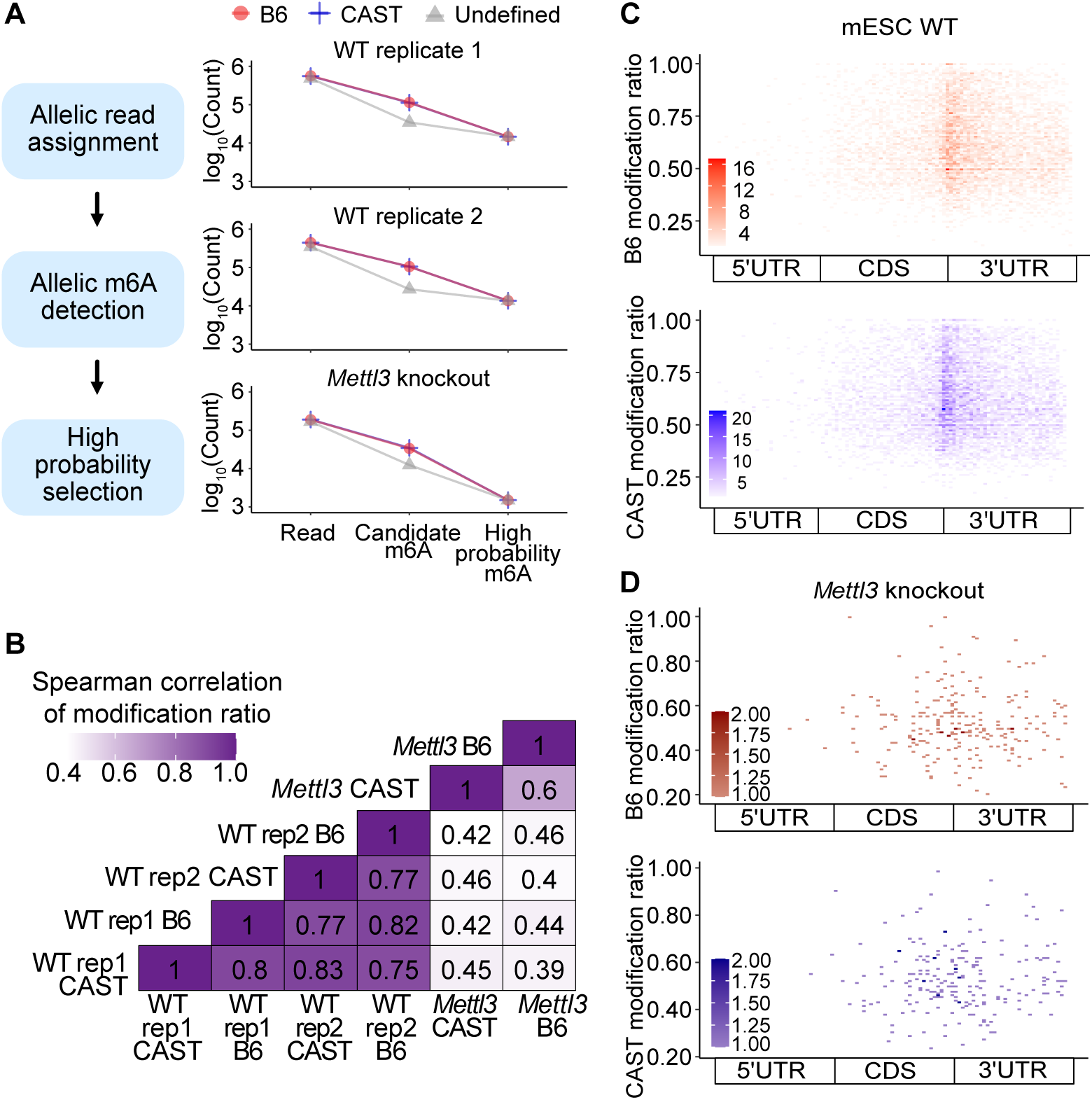
Comparative analysis of allelic modifications in wild-type and *Mettl3* knockout mESCs. A) Allelic impartiality while allelic read assignment and m6A detection. The left pane outlines the schematic of the data procedural steps for allele-specific m6A modifications analysis. The right panels display the counts for allelic reads, candidate m6A sites, and high probability m6A sites selected through our criteria (red circles, B6; blue plusses, CAST; and gray triangles, undefined group). The left top two plots show the counts from mESC wild-type (WT) replicates and the bottom plot exhibits the numbers from mESC *Mettl3* knockout. B) Spearman correlation of modification ratio between alleles from wild-type and *Mettl3* knockout cells (rep1: mESC WT replicate 1; rep2: mESC WT replicate 2; *Mettl3*: *Mettl3* knockout). C-D) Distribution of sites with high probability of m6A modification (prob > 0.85) is displayed in a metagene plot by calculating the relative positions of these sites within gene regions. The color scale represents the number of m6A sites with the given modification ratio inferred from reads assigned to either of the two alleles in mESC wild-type (C) or *Mettl3* knockout (D) cells (red, B6 allele; blue, CAST allele).

The modification ratios of the candidate m6A sites were highly correlated between replicates and demonstrated an even higher correlation within the same allele. Specifically, Spearman correlations within the same allele were 0.82 and 0.83 for the modification ratios of the B6 and CAST allele, respectively. Conversely, correlations between different alleles were slightly lower, with values of 0.77 (B6 replicate 1, CAST replicate 2) and 0.75 (CAST replicate 1 and B6 replicate 2). In contrast, modification ratios from *Mettl3* knockout cells exhibited significantly lower correlations, falling below 0.46 (Fig. 2B, Supplemental Fig. 3).

To identify allele-specific m6A modifications, we established a selection criterion centered on a site probability aggregated from all reads. Therefore, we focused on m6A sites that demonstrated a high probability of modification (>0.85) across reads. In mESC wild-type, an average of ∼7% of these candidate sites met our selection criteria. Notably, these m6A modification sites were predominantly localized at the junction between the coding region and the 3’ UTR, showing high modification ratios (with median ratios of 0.621 for B6 and 0.627 for CAST; Fig. 2A, C). In contrast, *Mettl3* knockout had only 0.8% allelic sites exhibiting high probabilities of m6A modification. Moreover, these sites demonstrated a wider dispersion across various transcript regions (Fig. 2A, D). Overall, these observations affirm the ability of our methodology in detecting allelic m6A modifications subsequent to the assignment of reads to alleles.

### Detection of sites with significant allele-specific m6A modification

The capability to accurately assign each RNA molecule to its allelic origin while concurrently identifying RNA modifications allows for the investigation of positions within mRNAs that exhibit differential modifications between alleles. While numerous statistical methods have been developed to identify allele-specific differences in gene expression phenotypes (DeVeale et al. 2012), these studies underscore the challenges inherent in this analysis, including a propensity for false positives when employing simple binomial tests to assess deviations from expected expression levels across the two alleles (Zitovsky and Love 2019; Mohammadi et al. 2017).

To address these challenges, we implemented a conservative strategy that leverages bootstrap sampling to quantify uncertainty in modification ratio estimates (Methods). This method enabled us to pinpoint mRNA positions showing significant allele-specific m6A modification (ASM) (Fig. 3A). Among detected 14,609 and 13,542 candidate m6A sites in the replicate experiments, we identified 57 ASM sites (FDR<0.1) with an average modification difference between the two alleles of 0.32.

**Figure. 3.**
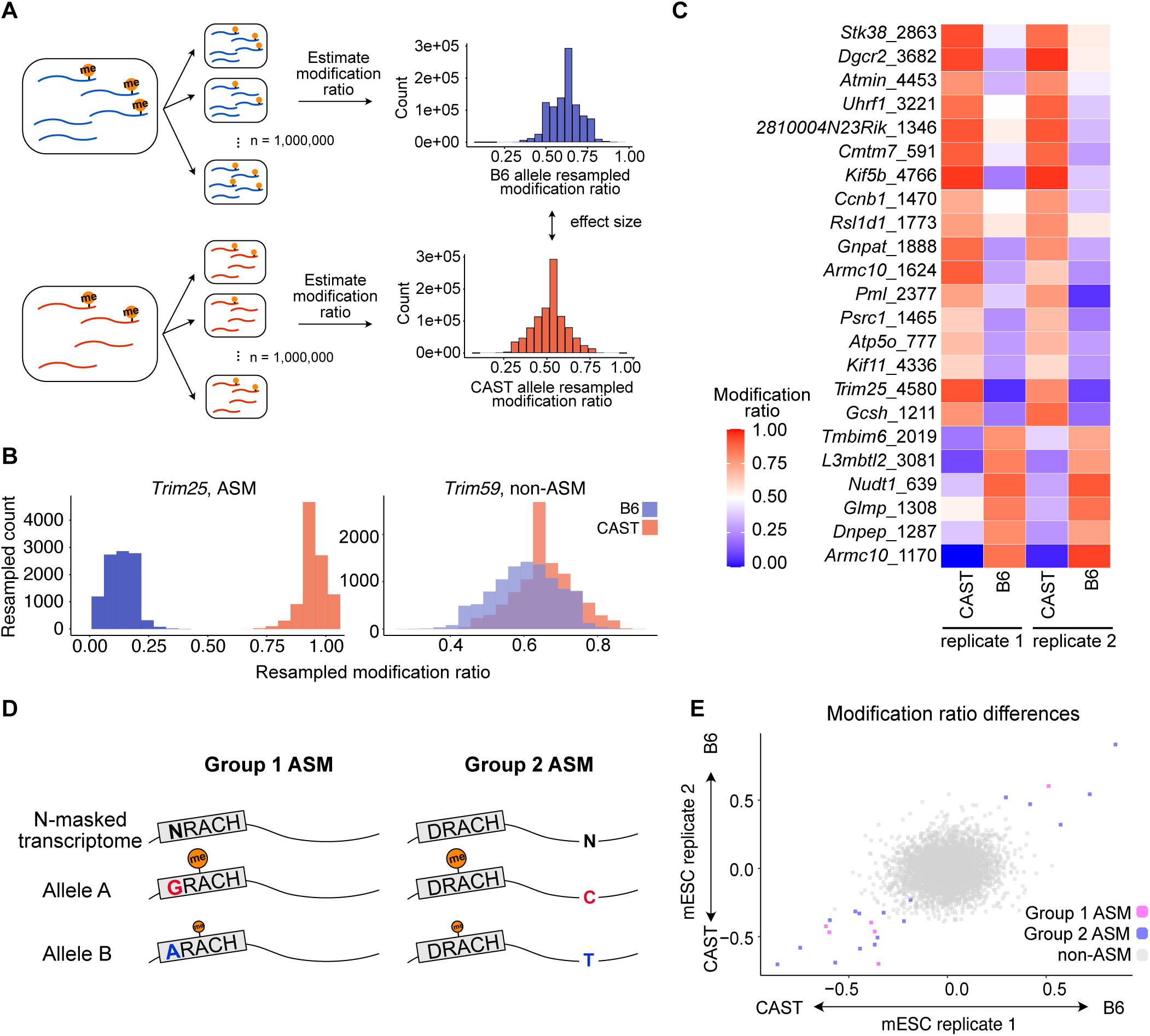
Identification and classification of ASM sites. A) Schematic of statistical procedure for ASM detection (Methods). Reads overlapping the site under consideration were resampled, and the modification ratio was estimated in each bootstrap sample. A statistically significant ASM site was defined as adjusted harmonic p-value (FDR < 0.1; Methods). B) An ASM site within *Trim25*, exhibits distinct modification ratio samples. Conversely, a non-ASM site within *Trim59* displays substantial overlap in modification ratios between bootstrap sampling distributions. C) Modification ratios of each allele across mESC wild-type replicates. Y-axis displays the name of the gene and m6A position in the transcript. D-E) ASM sites were classified into two groups. Group 1 is defined by genetic variants within the DRACH motif, and Group 2 is characterized by variants adjacent to or distal from the DRACH motif (D). The modification differences of the defined ASM were represented by color according to their classification (Group 1 in magenta, Group 2 in blue, and non-ASM in gray). Each axis is the modification ratio, where negative values denote CAST allele bias and positive values indicate B6 allele bias in m6A modification (E).

In allele-specific analysis, previous research revealed that events with larger effect sizes are more likely to be reproducible and biologically relevant (Castel et al. 2020; Mohammadi et al. 2017). Therefore, we repeated our statistical analyses using an effect size threshold of 0.1 corresponding to the inferred modification ratio difference between the two alleles (Methods). This analysis uncovered 23 sites across 22 genes indicating ASM. Notably, at these ASM sites, the distributions of the resampled modification ratios from the two alleles were consistently distinct and had large effect sizes, with a mean modification ratio difference of 0.48 (Fig. 3B, Supplemental Fig. 5A, Supplemental Table 3). We focused our detailed analyses on this subset of ASM sites.

One inherent limitation of Oxford Nanopore Technologies (ONT) sequencing lies in its limited sequencing depth, which constrains our ability to detect ASM in transcripts with low expression levels. Consistently, transcripts with statistically significant ASM sites had significantly greater RNA expression levels than those without, underscoring the dependence of ASM detection on transcript abundance (Supplemental Fig. 4).

The genes with ASM sites are distributed across chromosomes without any discernible location preferences (Supplemental Fig. 5B), and are associated with a wide range of functions (Fig. 3C). A particularly notable finding was the identification of two distinct sites of ASM on the *Armc10* transcript, which encodes a protein involved in mitochondrial dynamics (Chen et al. 2019; Serrat et al. 2014).

Moreover, our analysis identified six B6-biased ASM sites with a higher modification ratio on the B6 allele and 17 CAST-biased ASM sites with a higher ratio on the CAST allele. While the majority of ASM sites were located on 3’ UTRs, one B6-biased ASM (on *Dnpep*) and two CAST-biased ASMs (on *Gnpat*, and *Pml*) were found in coding regions, near the stop codons (Supplemental Fig. 5C).

Genome sequencing was used to identify genetic differences between the C57BL/6J (B6) and CAST/EiJ (CAST) inbred lines (Tsang et al. 2005; David J. Adams, Anthony G. Doran, Jingtao Lilue & Thomas M. Keane 2015), however, potential genotyping errors from these could result in erroneous ASM calls. To address this possibility, we verified the genomic DNA sequences near the m6A modification sites using Sanger sequencing (Supplemental Table 4, Methods). In six selected ASM sites, we confirmed annotated SNPs (*Nudt1*, D site), and the absence of unannotated genetic variants. These results indicate that the detected modifications are genuinely post-transcriptional and do not reflect genotyping errors.

Taken together, these findings highlight a key strength of our approach based on the ONT direct RNA sequencing technique, which enables the detection of m6A modifications at the individual molecule level, rather than relying on aggregate measurements.

### Genetic variants influence allele-specific m6A modification patterns

Having identified sites with ASM, we proceeded to examine the relationship between these sites and the genetic variants that differentiate the two alleles. We hypothesized that local genetic differences could influence methylation efficacy, leading to differential m6A deposition. Accordingly, we categorized ASM-biased sites into two groups based on the proximity of the nearest genetic variation to the canonical m6A methylation motif (DRACH) (Fig. 3D-E). Of the 23 sites identified, six had genetic variants located within the DRACH motif itself (Group 1), with three variants at the D position, one at R, and two at H (Fig. 4A). In total, 41 m6A sites had SNPs in D, R, or H positions, with 6 of these (14%) classified as ASM. These results suggest that SNPs within the DRACH motif are, as expected, more likely to lead to ASM. Furthermore, specific instances of the DRACH motifs are more likely to lead to modified adenines (Fig. 4B). In agreement with expectation, alleles for Group 1 ASM sites that exhibited higher modification ratios were more likely to match instances of the DRACH motif with higher propensity for modification (sole exception site on *Pml*).

**Figure. 4.**
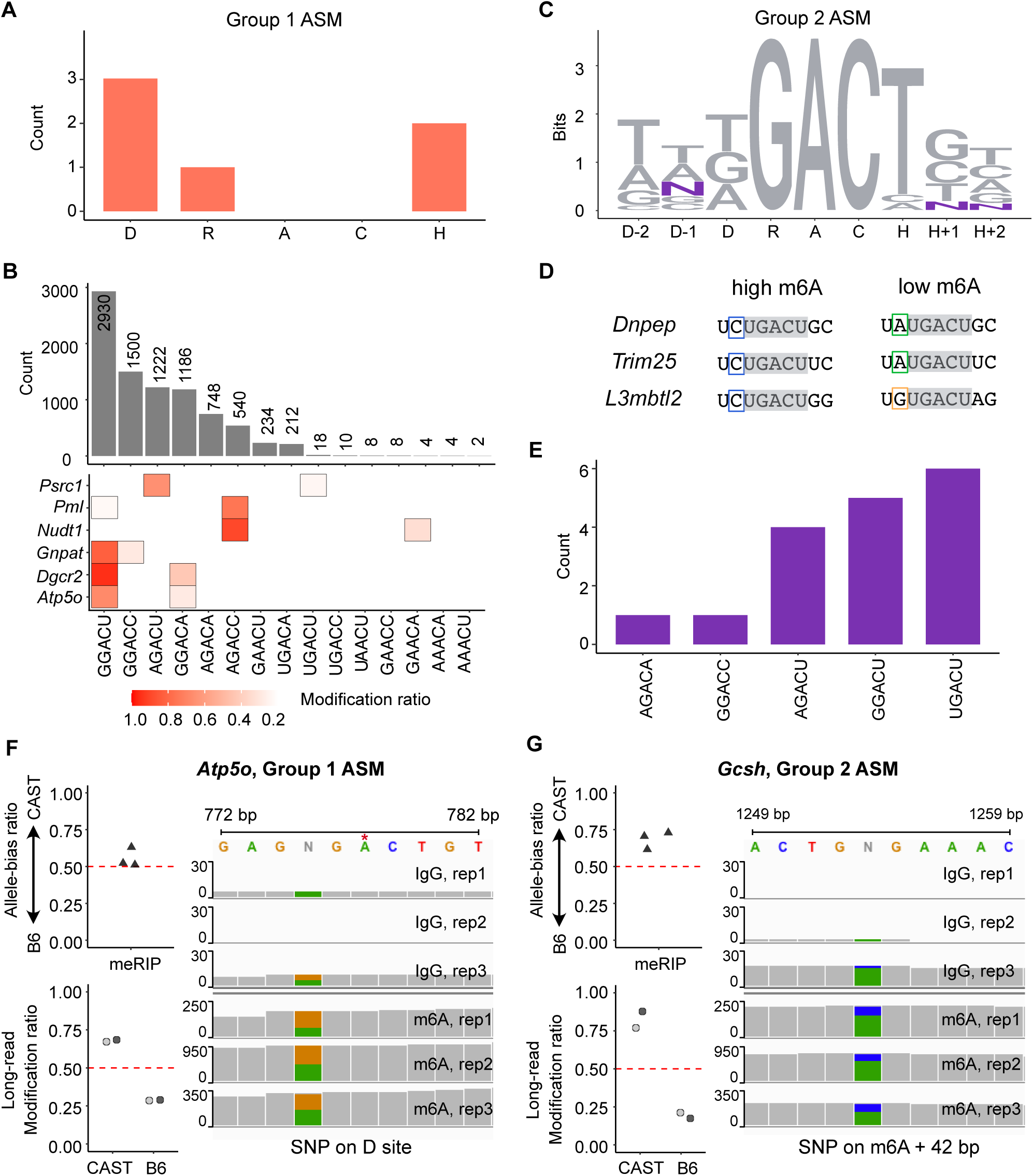
Characterization of ASM sites and orthogonal detection with MeRIP-seq. A) SNP distribution in Group 1 ASM. B) Motif frequencies and modification ratios of motif sequences. The top bar plot illustrates motif sequence frequencies in all m6A instances, while the bottom heatmap indicates modification ratios. The first row presents modification ratio of all instances and the following six rows represents modification ratio on each motif sequence differentiated by SNPs from two alleles of Group 1 ASM sites. C) Information content of the extended DRACH motif in 17 Group 2 ASM sites shown in DNA sequence. The D-1 site has three SNPs, while the H+1 and H-2 sites each have one SNP. D) Extended motif sequences where the D-1 site possesses SNPs. The gray box represents the DRACH motif, in which all three genes share the same sequence (UGACU) followed by U on the D-2 site. E) Motif prevalence in Group 2 ASM. The UGACU motifs are predominantly observed, contrasting with the common m6A motifs, which are typically represented by GGACU. F-G) Orthogonal detection of ASM through MeRIP-seq and long-read sequencing. On the top panel, points illustrate the allele-bias m6A ratio (proportion of reads from CAST allele) derived from three replicates of MeRIP-seq analysis. The Integrated Genome Viewer browser displays MeRIP-seq reads on SNPs adjacent to m6A sites, which correspond to the MeRIP-seq allele-bias ratio. The points in the bottom panel indicate the modification ratio of each allele from long-read sequencing, with gray color pairs representing data from two replicates. Two examples from *Apt5o*, Group 1 ASM (F) and *Gcsh*, Group 2 ASM (G) are shown.

Analysis of the remaining 17 sites (Group 2) revealed that five possessed genetic differences adjacent to the DRACH motif. Specifically, we found six SNPs near five m6A sites: *L3mbtl2* (D-1 and H+1); *Trim25* and *Dnpep* (D-1); *Atmin* and *Tmbim6* (H+2) (Fig. 4C). Intriguingly, all instances of SNPs at the D-1 position included a U at the D-2 position (UNUGACU). In this context, a cytosine at the D-1 site correlated with higher m6A levels (*Dnpep*, 0.768; *Trim25*, 0.855; *L3mbtl2*, 0.789) compared to an adenine or a guanine at the D-1 site (adenine on *Dnpep*, 0.324 and *Trim25*, 0.073; guanine on *L3mbtl2*, 0.157; Fig. 4D). This finding highlights the significant influence of nucleotides adjacent to the DRACH motif on m6A deposition, contingent upon their specific genetic context.

Among remaining Group 2 ASM sites, eight had a SNP within 100 base pairs of the modified adenine (*Stk38*, -64; *2810004N23Rik*, -51; *Cmtm7*, -12 bp; *Glmp*, +10 bp; *Rsl1d1*, +34 bp; *Gcsh*, +42 bp; Kif11, +59; Armc10 at 1170 position, +99). Despite the limitations imposed by the read length, short-read based m6A detection methods are theoretically capable of detecting SNPs within 50 to 100 bp of the methylated site (Dominissini et al. 2012; Chen et al. 2015). However, our method also identified four ASM sites that had no SNPs within this range hence highlighting the unique strength of long-read sequencing for ASM detection.

We also noticed that Group 2 ASMs were highly enriched for the UGACU motif sequence over the most commonly observed GAACU instance of the DRACH among m6A modified sites (Fig. 4E, p-val = 0.0084, Methods). This finding suggests that ASM may be more prevalent for specific motif sequences distinct from those typically seen in m6A modified sites. In summary, we uncovered differential m6A modification of alleles that may depend on genetic differences that are proximal to the DRACH motif as well as ASM sites which have no nearby genetic differences.

### Validation of ASM sites through orthogonal approaches

We next assessed the robustness of ASM detection by visualizing read pileups and conducting an orthogonal experimental method. Computational approaches to detect m6A modifications from direct RNA sequencing have been developed to leverage the increased propensity of base-calling errors around modified bases (Liu et al. 2019). To further validate sites that exhibited ASM using the supervised machine learning framework, we assessed the characteristic increase in errors surrounding each candidate site. We visualized sequencing reads that overlap ASM sites, enabling us to verify the expected enrichment of base-calling errors around sites with a higher modification ratio. The phenomenon was observed consistently across both replicates, characterized by a correspondence between base-calling errors and modification ratios (Supplemental Fig. 6).

An orthogonal experimental approach that can potentially detect transcript regions with ASM is MeRIP-seq (Cao et al. 2023). MeRIP-seq relies on antibodies to differentiate modified loci and utilizes short-read sequencing; thus, this strategy lacks single-molecule and single-nucleotide resolution. Nonetheless, we reasoned that some ASM sites would overlap MeRIP-seq peaks and provide additional experimental support for allelic bias.

Among the 23 ASM sites, we detected 20 in the MeRIP-seq. Only five out of 20 sites, which contain SNPs within or nearby the DRACH motif had sufficient coverage in our MeRIP-seq data (Supplemental Table 5, Method). The allele bias ratio measured from MeRIP-seq in these five sites demonstrated consistency with the allele bias detected by our approach (Supplemental Fig. 7). For example, *Atp5o* (Group 1 ASM) displayed allelic bias consistent with expectation in all three MeRIP-seq replicates (Fig. 4F). Another Group 2 ASM site, *Gcsh* (SNP at position 1,253, 42 bp downstream of methylation site), exhibited the same allele-bias pattern in both long-read sequencing and MeRIP-seq data (Fig. 4G). In short, while MeRIP-seq cannot capture all ASM sites detected by the long-read approach due to inherent limitations, we observed consistent allele bias in m6A patterns at five sites with sufficient read coverage.

The reliance of MeRIP-seq on short-read sequencing can lead to errors in allelic assignment, primarily due to dependence on a limited number of SNPs, which increases susceptibility to reference allele bias, genotyping errors, and systematic biases in library preparation. To assess potential genotyping errors, we examined 37 SNP sites within ASM genes using Sanger sequencing. Of these, 33 sites showed the expected genetic variants with strong peak signals, however, four sites (*Apt5o*, 776; *Psrc1*, 977; *Trim25*, 5007, and 5041) displayed nucleotides from only one allele. This finding suggests potential genotyping errors or limitations in our Sanger sequencing experiments. Importantly, these findings further underscore the challenges of accurate allelic detection especially for the short-read sequencing approach that rely on a one or few SNPs (Supplemental Table 6).

### Applicability of ONT direct RNA sequencing to detect ASM sites in human cells

The analytical and empirical workflow we developed to detect ASM sites is broadly applicable to any cell type with known genetic information. Given that systematic replication is essential to validate new approaches (Piccolo and Frampton 2016), we next replicated ASM detection using a lymphoblastoid cell line derived from a human with a well characterized genome. Specifically, we analyzed five replicates of ONT direct RNA sequencing data generated using the NA12878 cell line (Hansen 2016; Workman et al. 2018), assigned reads to their allelic origin, and quantified m6A modifications for each group of reads (Fig. 5A, Supplemental Table 7).

**Figure. 5.**
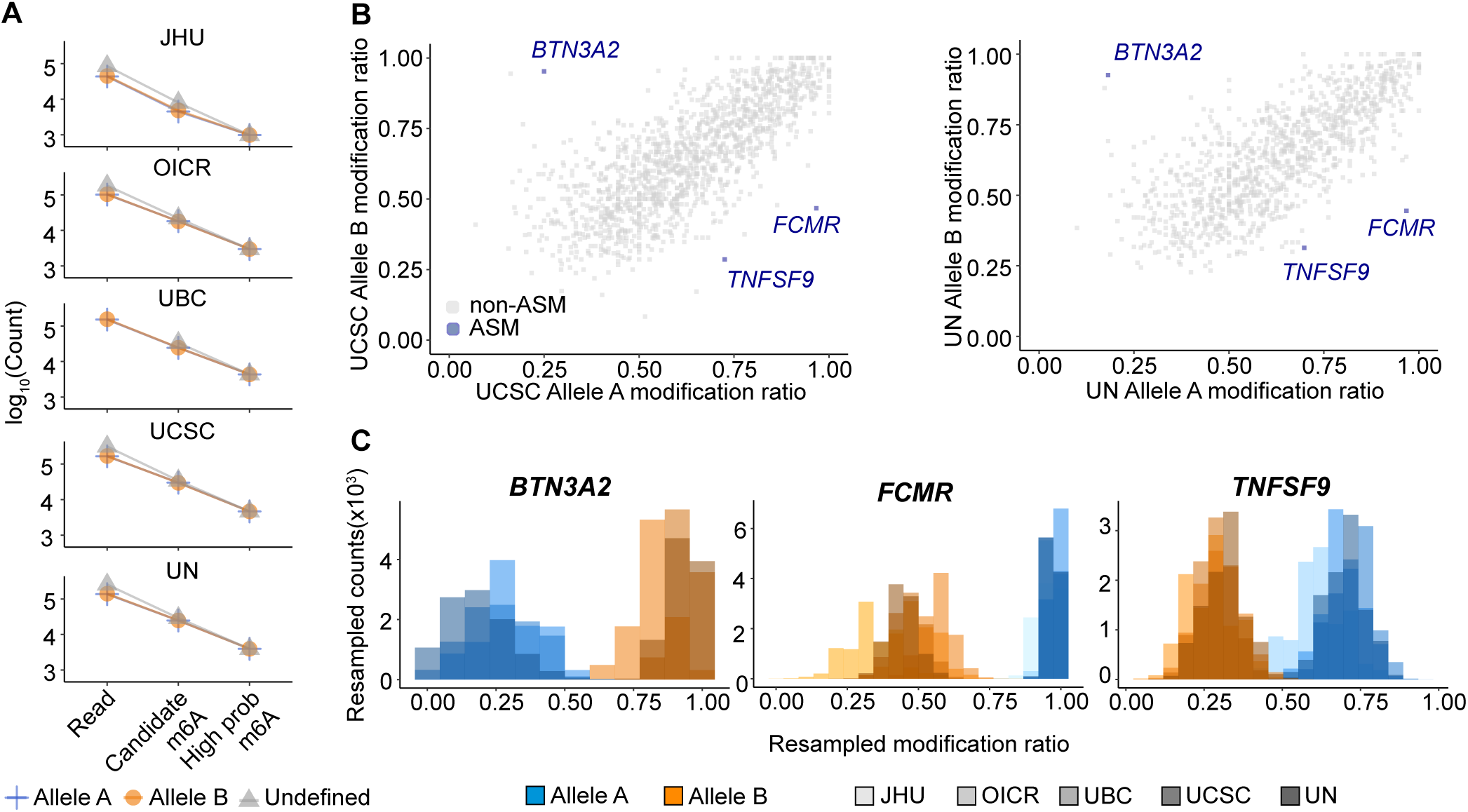
Reproducibility of ASM profiling procedure in human cells. A) Number of detected candidate m6A modification sites among five replicates (blue, Allele A; orange, Allele B; gray, undefined read classification). B) Modification ratios from each allele, including three ASM sites (blue) and non-ASM sites (gray) in UCSC (left) and UN (right) datasets, the highest depth datasets among five replicates. The X-axiswepresents the modification ratio of Allele A reads, while the Y-axis represents the modification ratio of Allele B reads. C) Resampled modification ratios from bootstrapping. Each color represents an allele (blue for Allele A, orange for Allele B), and the gray gradient indicates each replicate.

Unlike hybrid mouse ESCs, a typical human harbors many fewer heterozygous SNPs per transcript (Rozowsky et al. 2011; Workman et al. 2018). In our long-read sequencing analysis, among 21,569 mouse transcripts, 16,242 contain at least one heterozygous SNP in hybrid mESCs, while only 8,889 human transcripts contain such sites in NA12878. Notably, the transcriptome overall harbors nearly ten times fewer heterozygous SNPs in NA12878 compared to hybrid mESCs (210,004 in mESCs; 27,269 in NA12878; Supplemental Fig. 8). Hence, the percentage of long-reads that can be assigned to their allelic origin with high confidence is reduced (Supplemental Table 2, 4).

Despite having lower depth of sequencing and fewer informative SNPs per transcript, we identified three ASM sites with reproducible and large effect sizes. These three sites were found on the *BTN3A2* (FDR=0.006), *FCMR* (FDR=0.006), and *TNFSF9* (FDR=0.47) transcripts (Fig. 5B, Supplemental Fig. 9-10). *FCMR* contains a SNP on the R site of the DRACH motif. For the other two cases the closest SNP to the ASM site was found 86 (at position 3,175 in *BTN3A2* transcript) and 24 (at position 779 in *TNFSF9* transcript) nucleotides away from the methylation site. All three events were observed within the 3’ UTR and exhibited large differences in modification ratio between the two alleles (Fig. 5C; mean difference in modification ratio 0.657, *BTN3A2*; 0.500, *FCMR*; and 0.390, *TNFSF9*). *BTN3A2* plays a crucial role in T cell activation and proliferation (Vantourout et al. 2018; Kabelitz and Dechanet-Merville 2016), *FCMR*, which encodes the IgM Fc receptor, is vital for B cell activation and survival (Wang et al. 2016), and *TNFSF9*, a member of the TNF superfamily, enhances T cell responses by interacting with CD137 on activated T lymphocytes (Wang et al. 2016; Hashimoto 2021). Given that the NA12878 cell line is an Epstein-Barr virus (EBV)-transformed lymphoblastoid line, our findings reveal ASM sites within three transcripts related within the immune system. The applicability of our ONT direct RNA sequencing method for ASM detection in human cells supports the wide-ranging utility of our approach in any system with known genetic information.

### RNA abundance is higher for the allele with higher m6A modification ratio

The role of m6A RNA modification in transcription and translation has been extensively investigated and remains a topic of debate (Akhtar et al. 2021; Jain et al. 2023; Meyer 2019b). Allele-specific differences in m6A modification provides a powerful platform to assess their functional impact on expression dynamics as the genetic background, environmental factors and sample preparation are identical for the two alleles. Hence, we generated matched RNA-seq and ribosome profiling data in hybrid mESCs and leveraged existing measurements for the NA12878 cells (Methods) (Cenik et al. 2015). This data enabled us to determine the relative RNA expression and ribosome occupancy on each allele and correlate these with their m6A modification status.

In hybrid mESCs, transcripts harboring ASM sites demonstrated statistically significant RNA expression bias towards the allele with higher m6A modification. This pattern was consistent across both long-read and short-read sequencing methods (Fig. 6A; Binomial test p-value 0.004 and 0.011, respectively). Specifically, the mean proportion of RNA reads from the CAST allele for genes exhibiting CAST-biased ASM were 0.557 and 0.558 for long-read and short-read sequencing. Similarly, genes with B6-biased ASM had higher mean proportion of RNA reads from the B6 allele (0.460 and 0.398, respectively). These observations suggest that ASM is associated with allele-specific expression in the same allelic direction.

**Figure 6.**
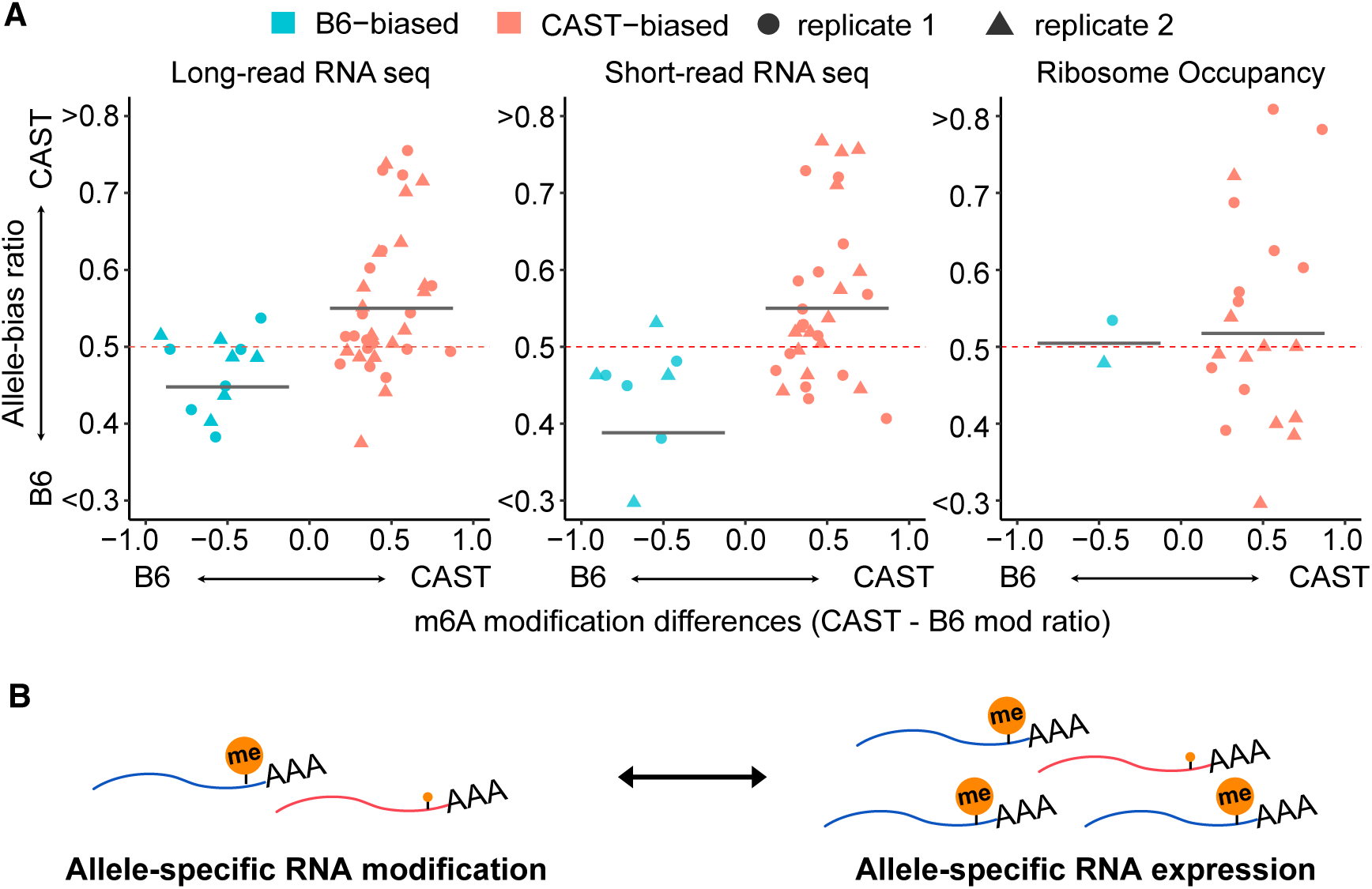
Effects of ASM on allele specific RNA expression and ribosome occupancy. A) Allele bias ratio of genes containing ASM sites (sky blue for B6 biased; pink for CAST biased). Different shapes represent the replicates (circle for replicate 1, triangle for replicate 2) The Y-axis displays the allele bias ratio obtained from long-read (left), short-read (middle) sequencing, and ribosome profiling (right). The X-axis shows the difference in m6A modification ratios between the two alleles (CAST - B6). The red dashed horizontal bar indicates allele-bias ratio 0.5, an allele-bias cutoff point. The gray bar represents the mean allele bias ratio for genes with B6 or CAST biased ASM sites. B) Model for regulation of ASM and allele-specific expression.

In NA12878 cells, the association of ASM and allele-specific RNA expression was similarly evident. *BTN3A2*, possessing Allele B biased methylation site, demonstrated a high proportion of RNA reads from Allele B (mean allelic ratios 0.851 and 0.704 for long-read and short-read sequencing). Similarly, *TNFSF9* and *FCMR*, with Allele A biased methylation sites, showed slightly elevated proportions of RNA reads from the Allele A (Supplemental Fig. 11A-B). These findings further support an association with ASM and allele-specific RNA expression (Fig. 6B).

Recent studies have postulated the role of m6A modification in regulating translation (Mao et al. 2019; Meyer 2019b; Jain et al. 2023). In light of this, we employed ribosome profiling to assess the impact of ASM on allele-specific translation. In particular, we analyzed allele-specific ribosome occupancy on genes with ASM in hybrid mESCs. We did not detect significant correlation between allele-specific ribosome occupancy and ASM (Fig. 6A; p-value, 0.83; Supplemental Fig. 11C). Collectively, our results indicate that alleles with higher m6A modification ratios are associated with increased RNA abundance but similar ribosome occupancy.

## DISCUSSION

In this study, we used ONT direct RNA sequencing as a new method to detect allele-specific m6A RNA modifications in both human and mouse cells. Notably, the long-reads generated by the ONT approach revealed ASM sites with no nearby genetic differences, suggesting that m6A modification on a given site may depend on factors beyond local sequence context. These ASM may potentially be governed from long-range interactions that modulate mRNA secondary structure, differences in allele-specific interactions with RNA-binding proteins or the local chromatin context of each allele (Huang et al. 2019; Deng et al. 2022; Berlivet et al. 2019).

The deposition of m6A modification on mRNA is dependent on the presence of a specific motif (DR**A**CH) surrounding the adenosine that is modified (Linder et al. 2015). Our analysis of ASM sites revealed that nucleotide identity of the positions that surround this canonical motif may also influence m6A deposition in particular contexts. Specifically, we found that alleles containing cytosine at the D-1 site followed by uracil at the D-2 site exhibit higher methylation levels (Supplemental Fig. 6).

A major strength of our approach over short-read based methods is its ability to cover many more informative SNPs to assign reads to their allelic origin (Supplemental Fig. 12, maximum SNP count per read, 12 with short-read; 78 with long-read in mESC). Hence, long-read technology has higher confidence in allelic assignment (Wu et al. 2023; Glinos et al. 2022). In contrast to short-read methods, which rely heavily on single SNPs within a read and are thus more susceptible to errors from misannotated variants, our approach minimizes the impact of incorrect or missing genetic variant annotations. Furthermore, in samples with less genetic variation, long-reads increase the chance of linking genetic variants that may be far away from the site of interest which would not be detectable by short-read based approaches.

A recent study leveraged previously generated MeRIP-seq data and claimed to detect numerous ASM sites (Cao et al. 2023). Their approach involved calculating p-values from Fisher’s exact test on tables of reads per kilobase of transcript per million mapped reads for each allele from the input control and IP. They interpreted the resulting p-values as evidence of ASM. However, this method is fundamentally flawed. Fisher’s exact test is specifically designed for categorical data, particularly in 2x2 contingency tables, and applying it to continuous data in this context is inappropriate. This misuse of the test raises serious concerns about the validity of their conclusions.

Furthermore, MeRIP-seq suffers from the additional limitations of antibody-based enrichment. Antibody-based approaches introduce specificity artifacts which result in variability in the number and location of peaks detected across experiments (Helm et al. 2019). Similarly, the immunoprecipitation step creates variable yields, limiting quantitative measurements among experiments (McIntyre et al. 2020). Therefore, the large number of sites reported by Cao et al. are inflated with a large number of false positives. In our study, we focused on large effect differences using a bootstrap resampling strategy and minimum effect size threshold to reduce statistical artifacts as previously recommended (Castel et al. 2020; Mohammadi et al. 2017). Consequently, the number of sites described here likely reflects the extent of allele-biased methylation more accurately.

To address the limitations of antibody-based detection of m6A modifications, recent work developed enzymatic and chemical approaches (Meyer 2019a; Song et al. 2021). However, the applicability of the enzymatic approach is currently restricted to a subset of m6A sites within DRACH motifs ending in ACA, constituting approximately 16% of total sites. While these advances are promising, they will likely be limited for allele-specific analysis due to the use of short-read sequencing (Garcia-Campos et al. 2019).

Allelic imbalances in m6A modification ratios between transcripts can potentially lead to allele-specific RNA expression and translation based on their impact on mRNA stability, transcription and translation efficiency (Mauer et al. 2017; Cesaro et al. 2023; Min et al. 2018) Specifically, m6A reader proteins such as YTHDC1 and YTHDC2, which interact with m6A sites on 3’ UTRs, are recognized for their role in enhancing mRNA stability and, consequently, increasing RNA abundance at steady-state (Lee et al. 2021; Wang et al. 2014). Our study revealed a positive relationship between ASM and allele-specific RNA expression. A potential mechanism explaining this association is the allele-specific association with m6A reader proteins that subsequently stabilize m6A-enriched mRNAs.

In contrast, we did not observe an association between ASM and allele-specific ribosome occupancy. Given that ribosome occupancy is a composite measurement of RNA expression and translation efficiency, this observation may indicate that alleles with higher modification ratios are less efficiently translated despite having higher steady-state RNA abundance. Such a mechanism would be in agreement with a previously proposed model of coupling between co-transcriptional m6A deposition and translation (Slobodin et al. 2017).

Our method has several important limitations. First, the supervised machine learning framework we adopted is predicated on the assumption that modifications occur exclusively within DRACH motifs. Consequently, our analysis does not account for genetic variations that alter the motif into sequences not matching the DRACH pattern, which are presumed to result in methylation loss. Second, the limitation in the number of reads generated by ONT direct RNA sequencing constrains our method’s ability to detect ASM sites in lowly expressed transcripts. Hence, ASM sites identified in this study occur in genes within the top 30th percentile of RNA expression (Supplemental Fig. 4).

In summary, we present a novel method for identifying allele-specific m6A modification using ONT direct RNA sequencing. Our analyses emphasize the benefits of long-read sequencing and direct detection of RNA modifications for ASM analysis. Future ASM studies are likely to extend the catalog of allelic variants that influence RNA modifications, and characterize the mechanisms leading to ASM and its functional consequences on gene expression.

## METHODS

### Cell culture

The C57BL/6J-CAST/EiJ F1 Hybrid mESCs were generously provided by Dr. David Spector (Balasooriya and Spector 2022). Cells were cultured in 2i medium, comprising Knockout DMEM (Gibco, Cat. No. 10829-018) supplemented with 15% Fetal Bovine Serum (FBS) (Millipore, Cat. No. ES-009-B), 1X Glutamine (Millipore, Cat. No. TMS-002-C), 1X non-essential amino acids (Millipore, Cat. No. TMS-001-C), 0.15 mM 2-Mercaptoethanol (Millipore, Cat. No. ES-007-E), 100 U/ml Penicillin-Streptomycin (Gibco, Cat. No.15140-122), 100 U/ml Lif (Millipore, Cat. No. ESG 1107), 1 μM PD0325901 (Sigma Aldrich, Cat. No. 444968), and 3 μM CHIR99021 (Sigma Aldrich, Cat. No. 361571). The culture plates were coated with 0.1% gelatin (Millipore, ES-006-B). The cells were cultured at 37 °C under 5% CO_2_ and passaged at a 70-80% sub-confluent state.

### Generation of *Mettl3* knockout mESCs

*Mettl3* knockout cells were generated by introducing Cas9/sgRNA ribonucleoprotein (RNP) complexes into mESCs via nucleofection (Kirton et al. 2013). The sgRNA was synthesized by Synthego (Supplemental Table 2). To form the RNP, 300 pmol of Cas9 protein (NEB, Cat. No. M0386M) and 600 pmol of sgRNA were incubated in Cas9 Buffer (150 mM KCl, 1 mM MgCl_2_, 10% v/v Glycerol, 20 mM HEPES–KOH [pH 7.5]) at room temperature for 30 minutes. Subsequently, 65 µL of 4D-Nucleofector X Solution was added to the RNP solution. Nucleofection was performed using the optimized protocol recommended by the manufacturer (SF Cell Line 4D-NucleofectorTM X Kit L). A cell pellet was collected from 2 x 10^6^ cells, resuspended in the RNP solution, and transferred into a 100 µL Nucleocuvette Vessel. Electroporation was carried out using the 4D-Nucleofector X Unit (Lonza) with the FF120 program. Post-nucleofection, cells were equilibrated at room temperature for 8 minutes, then transferred to a gelatin coated culture dish containing prewarmed 2i media. The cells were allowed to recover at 37 °C for 72 hours, followed by the isolation of single clones using serial dilution. The genomic DNA was isolated from cells grown from single clones and mutations were confirmed using the primers listed in Supplemental Table 2.

### Oxford Nanopore direct RNA sequencing

mESCs were grown in a 10 mm petri dish and collected from two different numbers of passages on separate days, considered as two biological replicates. The cells were lysed in TRIzol reagent (Zymo Research, Cat. No. R2050) and RNA was extracted according to the manufacturer’s instructions (Zymo Direct-zol RNA Kits, Cat. No. R2061). 5 μg of total RNA without poly(A) RNA isolation was used for direct RNA sequencing (Viscardi and Arribere 2022). The library was generated using the Oxford Nanopore Direct RNA Sequencing Kit (Nanopore Cat. No. SQK-RNA002) following the manufacturer’s protocol. The RNA sequencing from each RNA replicate was performed on four MinION MkIb with R9.4 flow cells (Oxford Nanopore Technologies Ltd.) with a 24-h runtime for each run.

### Human long-read data for method validation

We utilized published ONT direct RNA sequencing data from the human cell line NA12878 downloaded from https://github.com/nanopore-wgs-consortium/NA12878 (Hansen 2016; Workman et al. 2018). Five replicates were generated using RNA obtained from different institutes (JHU, Johns Hopkins University; OICR, Ontario Institute for Cancer Research; UBC, University of British Columbia; UCSC, University of California Santa Cruz; UN, University of Nottingham). Raw fast5 files were used directly for analysis (558,005 reads from JHU; 1,226,344 reads from OICR; 2,073,885 reads from UBC; 2,059,045 reads from UCSC; 1,686,124 reads from UN).

### m6A detection from ONT direct RNA sequencing

Following sequencing, we used Guppy v. 6.3.2 (quality score cutoff = 7) for base-calling from fast5 files and verified error rates with Pomoxis v0.3.15 (Buttler and Drown 2022). Reads were aligned to the transcriptome with minimap 2.1 (minimap2 ‘-ax map-ont). To reduce alignment biases, we used a transcriptome reference in which SNPs were masked with Ns as previously described (Ozadam et al. 2023) A mouse VCF was downloaded from the Mouse Genome Project (https://www.mousegenomes.org/), and the NA12878 VCF file was obtained from https://hgdownload.soe.ucsc.edu/gbdb/hg38/platinumGenomes/.

To identify m6A modifications, we first used Nanopolish v0.11.3 to generate an index with the ‘-- scale-events’ and ‘--signal-index’ options, aligning events to the N-masked transcriptome reference. Detection of m6A RNA modifications was conducted using m6Anet v-2.0.0 and a pretrained model (Hct116_RNA002) (Hendra et al. 2022). A minimum of 20 reads per site was required to call modification sites.

### Assignment of reads to their allelic origin

To assign aligned reads to their allelic origin, we first identified the positions on each read that correspond to a SNP, adjusting for any deletions and insertions in the read with respect to the reference. Initially, we selected reads that intersect a predefined number of SNPs. The number of heterozygous SNPs in mESC and NA12878 transcriptomes was 210,004, and 27,269, respectively (Supplemental Fig. 8). Consequently, we required at least three SNPs per read for mESCs, and one SNP for NA12878.

Then, we calculated the number of matches to each allele and defined a read-level allele-bias] ratio:

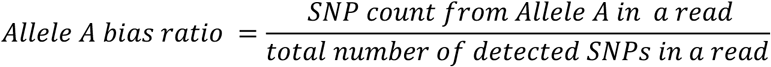

We assigned each read into one of three groups based on this ratio: allele A (bias ratio exceeding 0.7), allele B (bias ratio less than 0.3), and undefined (bias ratio between 0.3 and 0.7). The process was implemented in a python script that is provided on Github: allele_assignment.py. Subsequently, the three groups of reads underwent processing through m6Anet separately to predict m6A probability and modification ratios as described above.

### Identification of allele-specific m6A modifications

We first selected m6A sites with a high probability of modification (prob > 0.85) using all sequenced reads. When the SNPs on the motif convert DRACH motifs to non-DRACH motifs, we exclude them from the analysis because non-DRACH motifs are by definition assumed to be unmethylated (Hendra et al. 2022). We select the case in which SNPs on the motif do not make changes in the DRACH motif. In mESCs, 29 out of 178 in replicate 1 and 29 out of 145 in replicate 2 were further evaluated, as the SNP overlapping the motif led to different instances of the DRACH sequence (Supplemental Fig. 13). In NA12878, the corresponding numbers were 1 in 5 (JHU), 1 in 11 (OICR), 1 in 21 (UBC), 2 in 24 (UCSC), 2 in 22 (UN).

Given that m6Anet sets a threshold of 20 reads for determining modification sites and ratios, sites with fewer than 20 reads in one of the allelic groups are excluded during the detection phase. This results in a discrepancy between the number of reads assigned to alleles A, B, the undefined category, and the total count of reads. To correct this disparity and obtain an accurate modification ratio for these sites, we initially identified modification sites in transcripts that harbor at least one heterozygous SNP and at least 20 reads assigned to one of the alleles. If both alleles had more than 20 reads, the modification ratios were used directly as calculated by m6Anet (63% of total instance). However, when one allele has read counts less than 20, we recalculated the modification ratio leveraging modification information from all reads, without distinguishing the two alleles. Without loss of generality, let’s assume that allele A had fewer than 20 reads assigned and hence was not considered by m6Anet. In this case, we first calculated its read count by following:

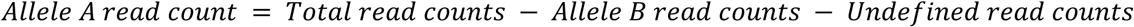

If at least 10 reads were assigned to allele A, we retained this site for further analysis and recalculated the modification ratio of allele A using the following formula:

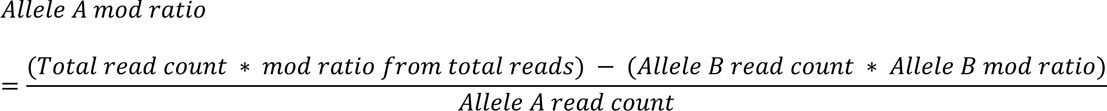

See Supplemental Fig. 14 for a schematic description of this procedure.

We identified statistically significant ASM sites using a bootstrapping-based statistical test. First, for each allele, methylated read counts were derived by multiplying modification ratios with total read numbers. We then sampled the number of methylated reads for each allele with replacement and calculated the difference between the modification ratios using the resampled read counts (McLachlan and Rathnayake 2014; Banjanovic and Osborne 2016). This resampling procedure was repeated 1,000,000 times and a one-sided p-value was calculated by using a effect size threshold (*T*) of 0 or 0.1 as follows:

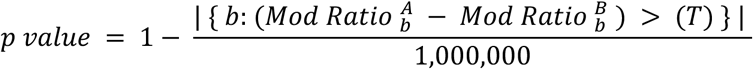

where, 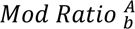 represents the bootstrap resampling value for the allele with the higher observed m6A modification ratio. An aggregate p-value was calculated by combining the p-values from each replicate using the harmonic mean method (Wilson 2019). False Discovery Rate (FDR) was calculated by the Benjamini & Hochberg method (Yoav Benjamini 1995). Finally, statistically significant ASM sites were defined if adjusted harmonic mean p-values (FDR) were below 0.1. For instance, with an effect size threshold of 0.1 (T = 0.1), if none of the randomizations exceed this cutoff, it would suggest that the modification ratios of the two alleles from resampled reads are highly similar. Consequently, the probability of this site being an ASM would be very low, corresponding to a p-value of 1.

To assess the significance of UGACU being the most common DRACH variant among Group 2 ASM sites, we randomly resampled 17 motifs 10,000 times. For the resampling, we used the observed frequency of each of the 15 instances of the DRACH motif among sites with a high probability of modification. In these random samples, only 84 instances had UGACU as the most frequent motif hence corresponding to a p-value of 0.0084.

### mESC MeRIP-seq experiments and analyses

MeRIP-seq libraries were prepared with EpiNext CUT&RUN RNA m6A-Seq Kit (EpiGentek). The three replicates of mESCs were collected from different numbers of passages on separate dates. The total RNA was extracted with Direct-zol RNA Purification Kits (Zymo Research, Cat. No. R2050). 7 µg of total RNA was subjected to immunoprecipitation with an m6A antibody (P9016, EpiGentek, 1:100 dilution) and digested with cleavage enzyme on beads. The beads were then washed three times with a wash buffer and protein digestion buffer, and RNA was eluted in 13 µl of the elution buffer. The sequencing libraries were generated using the Diagenode small RNA sequencing kit following the manufacturer’s protocol (Diagenode, Cat. No. C05030001). The libraries were sequenced on a NovaSeq6000 system (Illumina).

Adaptor sequences were trimmed from raw reads with cutadapt v4.7 (Martin 2011) using following parameters: -a AAAAAAAAAACAAAAAAAAAA -G ^TTTTTTTTTGTTTTTTTTTT -AAGATCGGAAGAGCGTCGTGTAGGGAAAGAGTGT -n 2 --overlap=4 --trimmed-only --maximum-length=150 --minimum-length=31 --quality-cutoff=28. Trimed reads were aligned to the N-masked mouse transcriptome with STAR v2.7.10b (Dobin et al. 2013). Reads with low mapping quality were discarded (mapping quality less than 10) and indexed with samtools v1.15.1 (Danecek et al. 2021; Bonfield et al. 2021).

To compute the allele bias ratio, we counted the number of allelic reads that harbor at least one SNP within 100 bp of the ASM sites. Out of 23 ASM sites, four did not have genetic differences within 100 base pairs of the methylated position and 15 had fewer than 40 reads across the three replicates (Supplemental Table 5). Allele bias for the remaining five sites were calculated as:

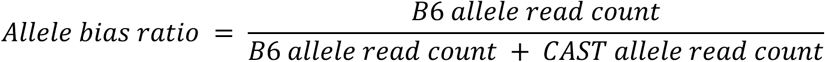

### mESC RNA-seq and ribosome profiling library preparation

Five million mESCs were scraped and transferred to 1.5 mL tubes in lysis buffer (20 mM Tris HCl pH 7.4, 150 mM NaCl, 5 mM MgCl_2_, 1 mM DTT, 100 µg/mL Cycloheximide, 1% Triton-X). All experiments were done in two replicates. Cells were lysed on ice by pipetting up and down ∼5 times every 5 min for a total of 10 min. The lysate was clarified by centrifugation at 1300 x g for 10 min at 4 °C. Ten percent of the clarified lysate by volume was reserved for RNA extraction using Direct-zol RNA Purification Kits (Zymo Research, Cat. No. R2050). The RNA-seq libraries were prepared using the NEBNext Ultra™ II RNA Library Prep Kit for Illumina following manufacturer’s protocol by Novagene.

The rest of the supernatant was immediately processed for ribosome profiling. Briefly, 7 μL of RNaseI (Invitrogen, Cat. No. AM2249) was added to the clarified lysates and digestion was carried out for 1 h at 4 °C. The digestion was stopped with ribonucleoside vanadyl complex (NEB, Cat. No. S1402S) at a final concentration of 20 mM. Digested lysates were layered on a sucrose cushion (20 mM Tris HCl pH 7.4, 150 mM NaCl, 5 mM MgCl_2_, 34% sucrose, 1 mM DTT) and the ribosomes were pelleted by centrifugation in a SW 41 Ti rotor (Beckman Coulter) at 38,000 rpm for 2.5 h at 4°C. RNA was isolated with the RNeasy Mini RNA Kit (Qiagen, Cat. No. 74104) and size-selected by running 5 µg of each sample on a 15% polyacrylamide TBE-UREA gel. The 21– 34 nt RNA fragments were excised and extracted by crushing the gel fragment in 400 µL of RNA extraction buffer (300 mM sodium acetate pH 5.5, 1 mM EDTA, 0.25% SDS) followed by a 30 min incubation on ice and an overnight incubation at room temperature. The sample was passed through a Spin X filter (Corning, Cat. No. 8160) and the flowthrough was ethanol precipitated in the presence of 5 mM MgCl_2_ and 1 µL GlycoBlue (Invitrogen, Cat. No. AM9516). The RNA pellet was resuspended in 10 µL of RNase-free water and immediately processed for library preparation.

For ribosome profiling library preparation, the D-Plex Small RNA-seq kit (Diagenode, Cat. No. C05030001) was used with slight modifications. The dephosphorylation reaction was supplemented with 0.5 μl T4 PNK (NEB, Cat. No. M0201S), and the reaction was incubated for 25 minutes. Subsequently, the complementary DNA (cDNA) was amplified for 12 PCR cycles. We used AMPure XP bead cleanup (1.8X), followed by size selection using 3% agarose, dye-free gel cassettes with internal standards (Sage Science, Cat. No. BDQ3010) on the BluePippin platform. Sequencing was performed on a Novaseq 6000 platform.

### Read processing of RNA-seq and ribosome profiling

For mESC, RNA-seq and Ribo-seq data were processed using RiboFlow v0.0.1 (Ozadam et al. 2020). For the Ribo-seq library, Unique Molecular Identifier (UMI) sequences were isolated employing the following parameters: “umi_tools extract -p ‘^(?P<UMI_1>.{12})(?P<DISCARD_1>.{4}).+$’ --extract-method=regex”. Subsequently, reads underwent clipping with the parameters “-a AAAAAAAAAACAAAAAAAAAA --overlap=4 --trimmed-only”. Trimmed reads were then filtered by alignment to mouse rRNA and tRNA sequences with bowtie2 version 7.3.0 and utilizing unaligned reads for subsequent alignment to the N-masked transcriptome (Langmead and Salzberg 2012). Following transcriptome alignment, reads with mapping quality greater than two were preserved and deduplicated utilizing UMI-tools directional adjacency method with the parameter “--read-length” (Smith et al. 2017).

In mESC RNA-seq analysis, we clipped NEB Read adaptors using cutadapt, v1.18 (Martin 2011) with following parameters: “-a AGATCGGAAGAGCACACGTCTGAACTCCAGTCA -A AGATCGGAAGAGCGTCGTGTAGGGAAAGAGTGT -O 8 -m 20 --cores=8”. The reads were aligned to the N-masked transcriptome using bowtie2 (Langmead and Salzberg 2012). The read count for RNA-seq and Ribo-seq were obtained from .ribo files with RiboR (Ozadam et al. 2020).

For the NA12878 sample, we analyzed RNA-seq and ribosome profiling data (NCBI Gene Expression Omnibus (GEO) under accession number GSE65912) based on the study by Cenik et al. (Cenik et al. 2015). We processed reads from both methods using cutadapt v1.18 with the parameters “-a AGATCGGAAGAGCACACGTCTGAACTCCAGTCA -AAGATCGGAAGAGCGTCGTGTAGGGAAAGAGTGT -O 8 -m 20” for trimming. The trimmed reads were then filtered by aligning to human rRNA and tRNA sequences with bowtie2 v7.3.0. Reads that did not align were subsequently mapped to the N-masked human transcriptome.

### Allele-specific RNA expression and ribosome occupancy analysis

Utilizing the aligned BAM files obtained from RNA-seq and Ribo-seq, ASE counts were acquired using GATK (version 3.8.1) ASEReadCounter (McKenna et al. 2010). The fraction of reads corresponding to the two alleles was calculated for all loci. After normalization by count per million reads, ASE scores were computed by dividing the read count from a certain allele to the sum of the read counts from both alleles (Castel et al. 2015; Liu et al. 2018).

To compare the allele-specific RNA expression and ribosome occupancy ratio in genes which have ASM, we obtained allele bias ratio to the same allele (e.g., allele A) which showed ASM (e.g., allele A bias methylation).

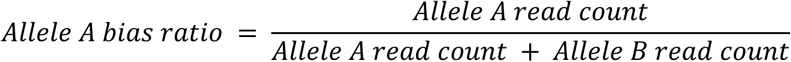

To quantify the relationship between the allele-bias ratio from long-read sequencing and short-read sequencing, we calculate the weighted Spearman correlation using long-read sequencing read counts as weights. The correlation was calculated by using R package “boot”, version 1.3-30.

### Genomic DNA extraction and PCR for genetic variant verification

Genomic DNA from mESCs was extracted using the Quick-DNA Miniprep Plus Kit (Zymo, Cat. No. D4068) following the manufacturer’s protocol. 17 primer pairs were designed to detect genetic variants at genomic ASM and SNP sites (Supplemental Table 8). The target regions were amplified by PCR using Q5 High-Fidelity DNA Polymerase (NEB, Cat. No. M0491S). The thermal cycling conditions were set as follows: initial denaturation at 95°C for 90 seconds, followed by 32 cycles of denaturation at 95°C for 10 seconds, annealing at the (primer melting temperature −2°C) for 15 seconds, extension at 72°C for 20-40 seconds, and a final extension at 72°C for 5 minutes. The resulting PCR products were purified using the NucleoSpin Gel and PCR Clean-up Kit (Takara, Cat. No. 740609.250) and sequenced by Sanger sequencing (ACGT, Inc. DNA sequencing service).

## DATA ACCESS

All mESC short-read sequencing data sets presented in this paper have been deposited in the Sequence Read Archive under BioProject accession number PRJNA1071025 (SRP486746). The ONT direct RNA sequencing data is available on Zenodo under the following record numbers: mESC replicate 1 (10815502, 13255832, 13256383), mESC replicate 2 (13257639, 13259594, 13273847, 13275906, 13278114, 13277067), and Mettl3 knockout cells (13257082).

All custom scripts used to perform bioinformatics analyses available on Github: https://github.com/DayeaPark/Allele-specific-m6A-modification.git

## ACKNOWLEDGMENTS

We thank Dr. David Spector for kindly providing hybrid mESCs. We appreciate the insightful comments provided by Dr. Ian Hoskins on the manuscript. This work was supported by National Institutes of Health grants [HD110096, GM150667], as well as a Welch Foundation grant [F-2027-20230405] (C.C.). Figures were generated using BioRender.com under a publication license (JO26GB978U). All original text in this paper was authored by the researchers. Additionally, we acknowledge the assistance of a Large Language Model (OpenAI, ChatGPT v3.5) for suggesting edits aimed at improving clarity and grammar.

