## Supplemental Figures for "Long-read RNA sequencing reveals allele-specific N^6^-methyladenosine modifications"

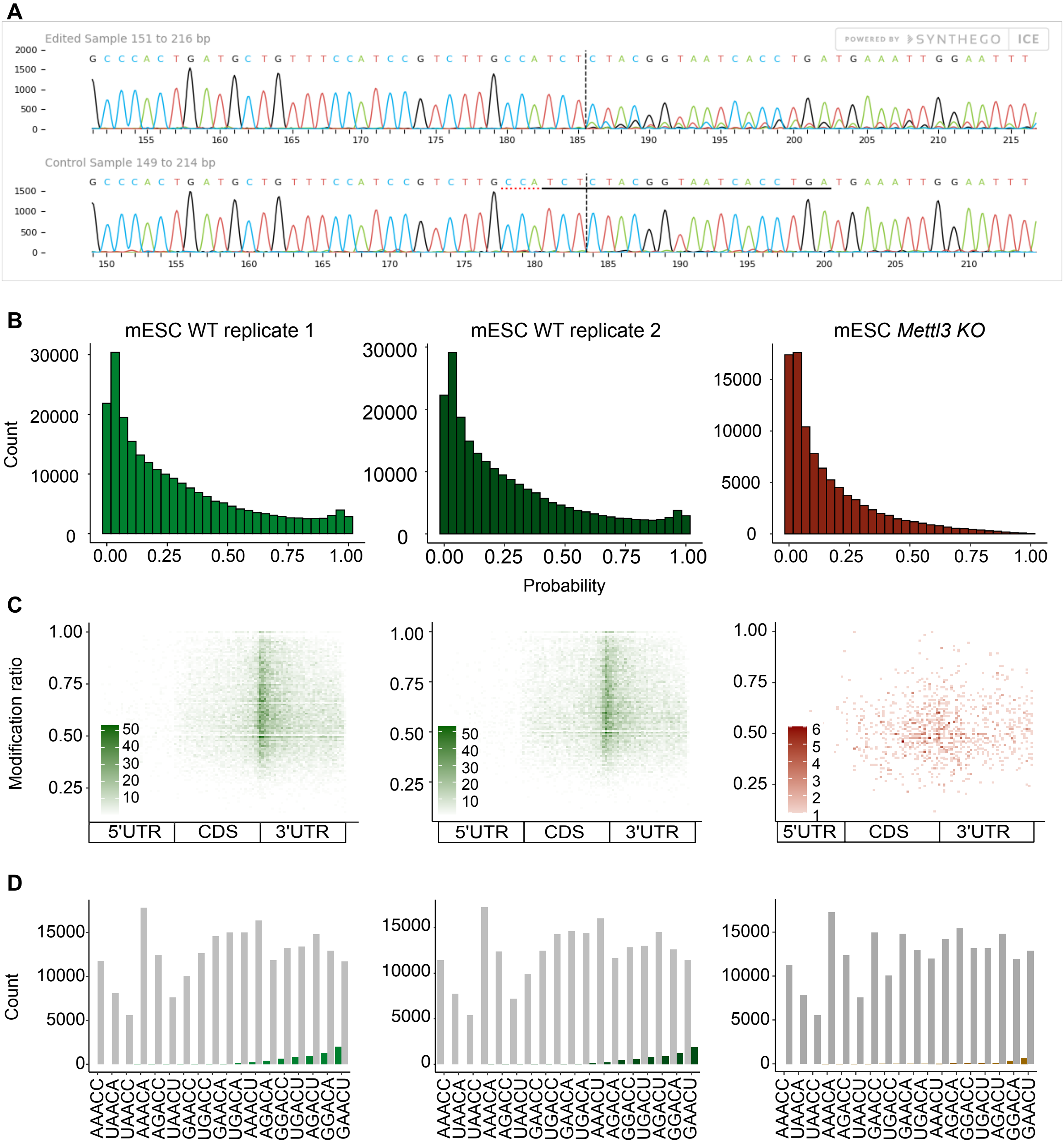


**Supplemental Figure. 1 | Mettl3 knockout versus wild–type direct RNA sequencing quality**

A) Mettl3 knockout target site located in exon 2. The image is generated using Synthego Inference of CRISPR edits. The interference was started from the 180 bp region where the sgRNA target site was located. The upper layer represents Sanger sequencing signal for Mettl3 knockout, and the bottom layer represents sequencing signal from wild-type. The PAM sequence is underlined with a red dash line, and the target site is underlined with a black line. B) The distribution of modification probabilities from two mESC WT replicates (green) and Mettl3 knockout (red). C) Modification ratios and locations of m6A sites with high probability ($p > 0.85$) of being modified were plotted. The relative m6A positions within the transcript body were determined. The color scale indicates the number of sites. D) Comparison of motif sequence preference before (gray) and after (colored) selection of sites with a high-probability ratio.

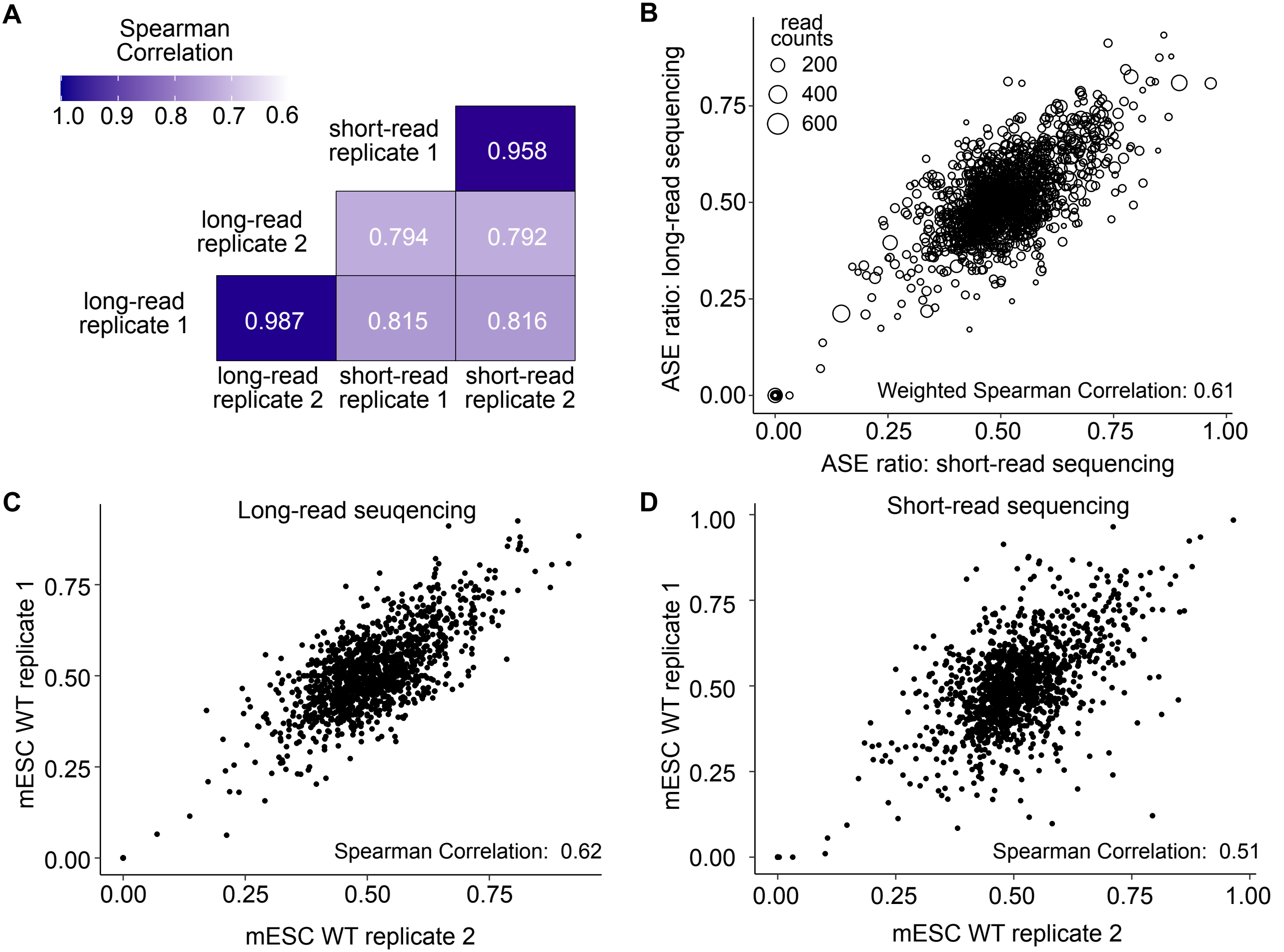


**Supplemental Figure. 2 | ASE ratio correlation between long-read and short-read sequencing.**

A Per transcript RNA expression was calculated by counts per million normalization and compared. B) Transcript-level Allele-Specific Expression (ASE) ratios were calculated and plotted using CAST allele ratio (CAST read counts) / (CAST read counts + B6 read counts). The correlation between estimated ASE ratio was calculated between long-read and short-read sequencing using the mean of the replicates weighted by read counts from long-read sequencing (depicted with the size of the circles). The correlation between replicates from each method exhibited similar correlation levels (C, D).

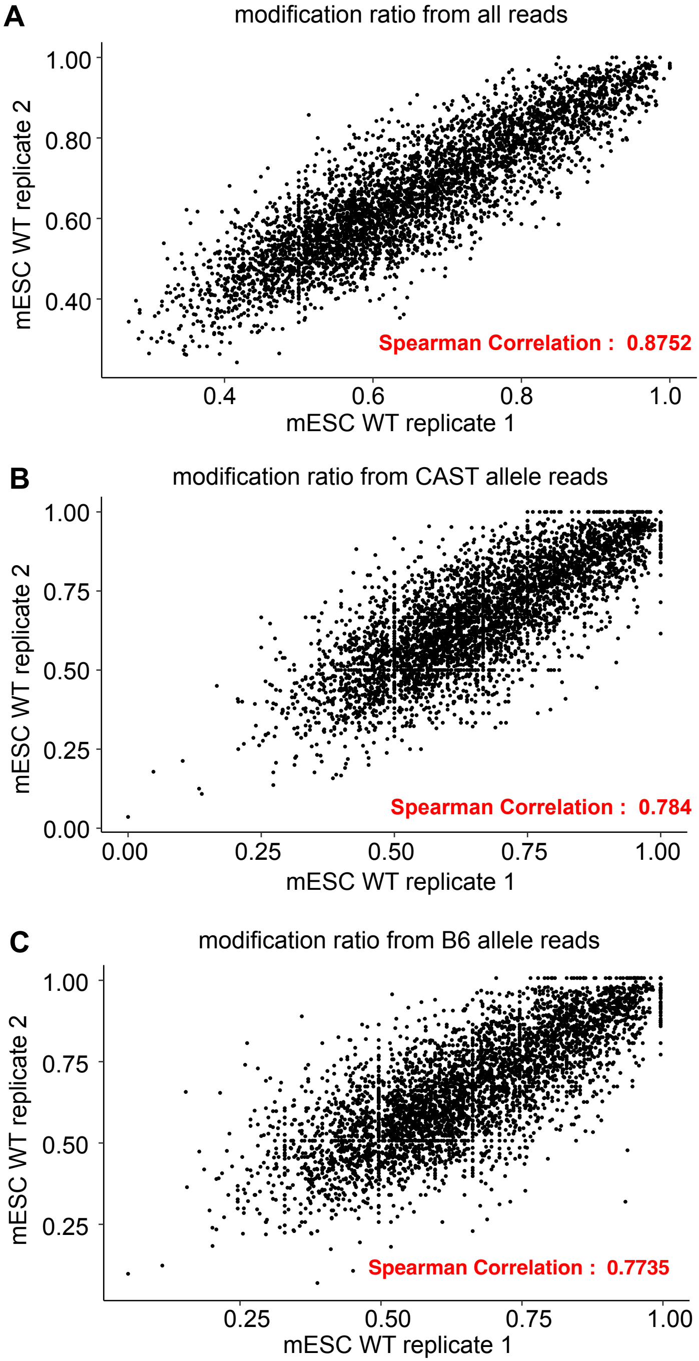


**Supplemental Figure. 3 | Correlations of modification ratios between replicates**

Spearman correlations were measured for modification ratios calculated using all reads (A), CAST allele reads (B), and B6 allele reads (C).

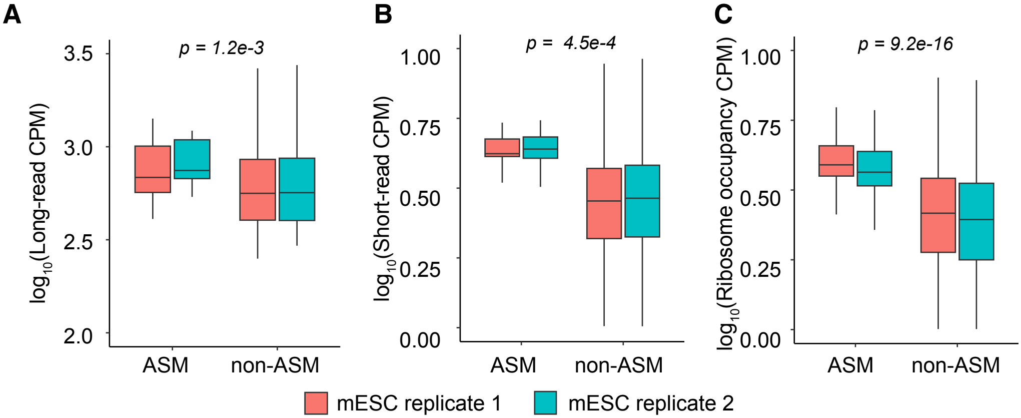


**Supplemental Figure. 4 | Detection of ASM is biased towards more highly expressed genes**

Comparison of RNA expression and ribosome occupancy of transcripts harboring sites with ASM or without ASM. RNA expression and ribosome occupancy were calculated using counts per million reads (CPM) from long-read sequencing (A), short-read sequencing (B), and ribosome profiling data (C). The boxplot compared RNA expression level and ribosome occupancies of the transcripts which contain ASM or not. P-values were obtained using a two-sided Wilcoxon Rank-Sum Test.

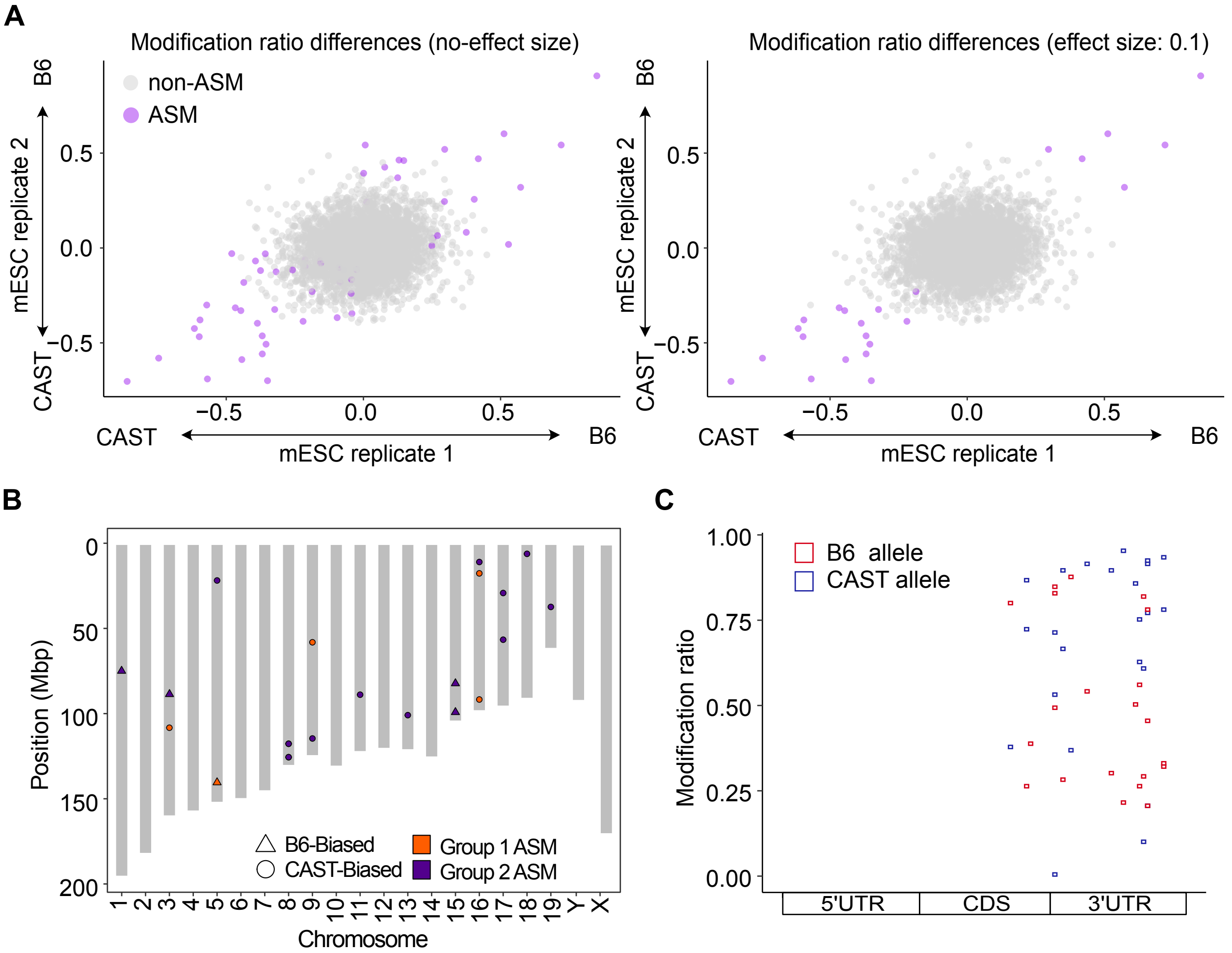


**Supplemental Figure. 5 | Characterization of ASM sites**

A) Comparison of ASM determinations methods using no-effect size (left) or 0.1 (right; Methods). Non-ASM sites are colored in gray and ASM sites in purple. B) Chromosomal distribution of ASM sites in male hybrid mESCs. C). Gene region distribution and allelic modification ratios on ASM sites. The relative m6A positions within the transcript body were determined. Colors signify the modification ratios for the B6 (orange) and CAST (blue) alleles, highlighting the allelic differences in modification.

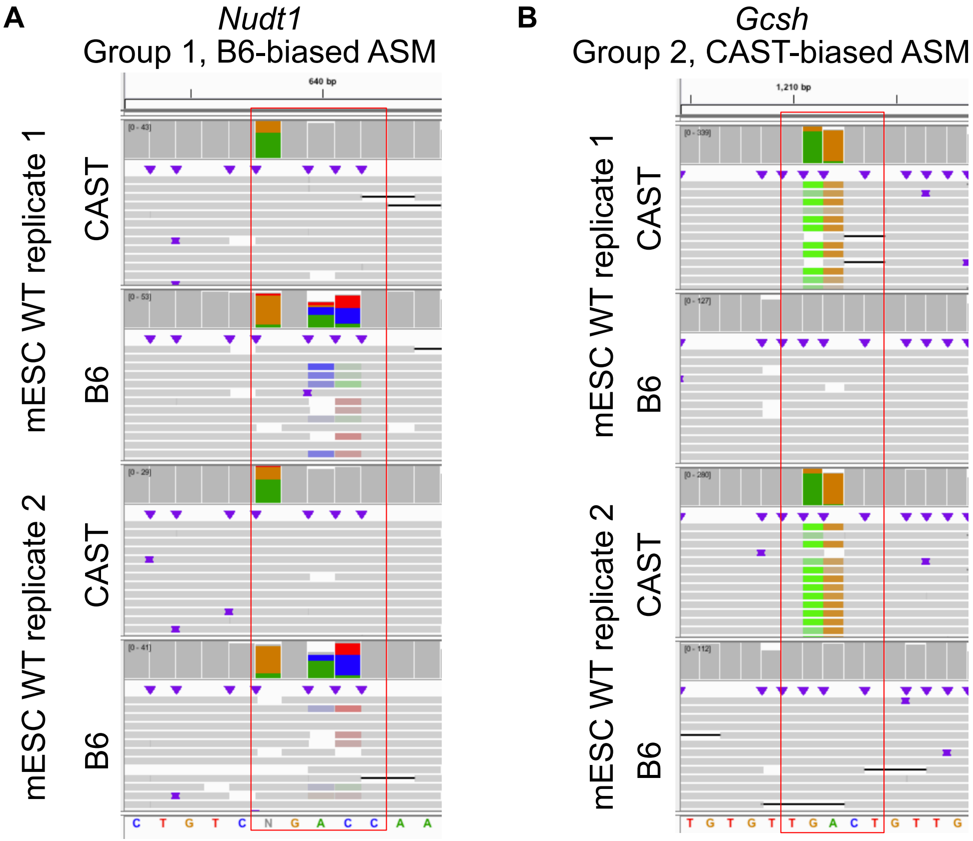


**Supplemental Figure. 6 | Visual detection of ASM from long-read sequencing**

Visualizing base-calling errors in the Integrated Genome Viewer browser confirm the presence of ASM sites. A) Nudt1 was identified as B6-biased ASM, showing a high-error rate in reads from B6 allele. B) Gcsh, classified under Group 2 CAST-biased ASM, demonstrates a high frequent error on reads from CAST allele. The red box represents the DRACH motif.

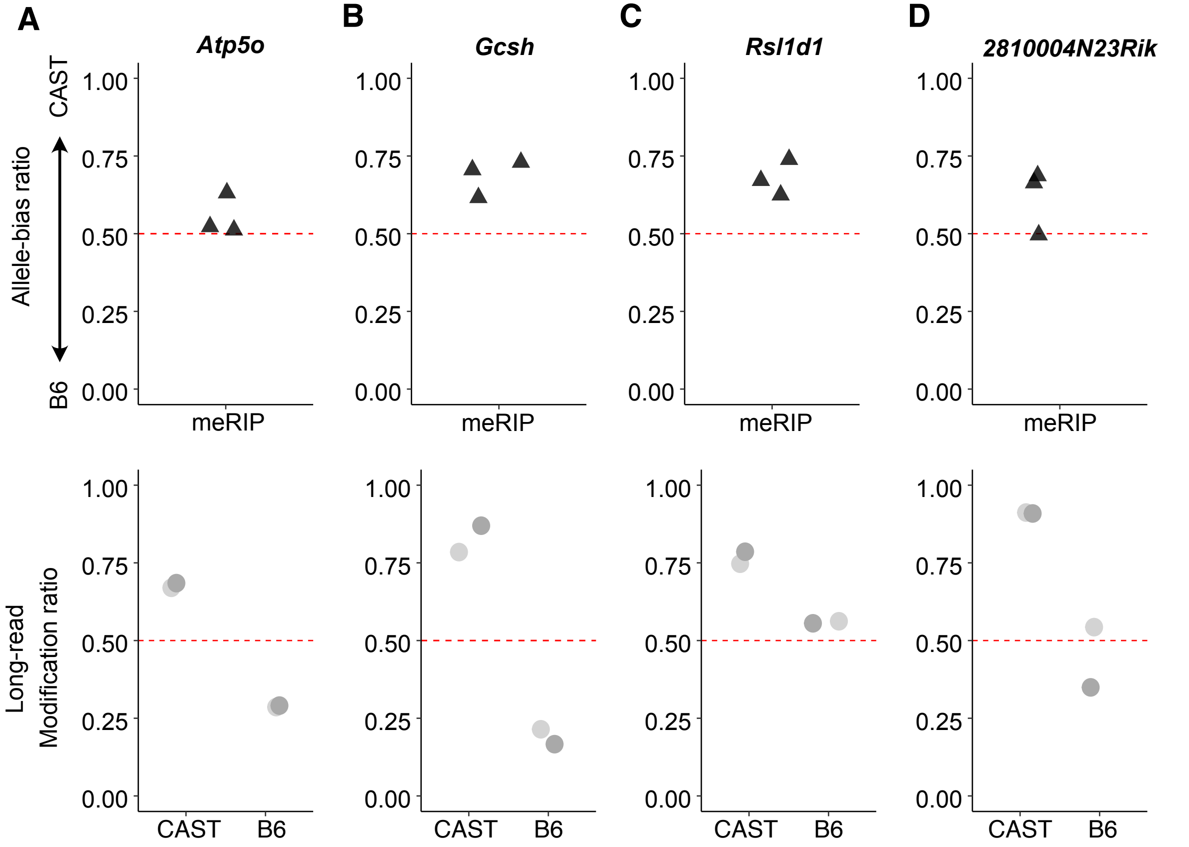


**Supplemental Figure. 7 | Comparison of allele bias ratio from MeRIP-seq and allelic modification ratio from long-read sequencing.**

Five ASM sites (Group 1 (A) and Group 2 (B, C, D)) were detected by both MeRIP-seq and long-read sequencing. Top panels represent allele-bias ratio (proportion of reads from CAST allele) from MeRIP-seq; therefore, a high ratio implies allele bias toward CAST allele and low ratio indicates a bias toward B6 allele. The points in the bottom plot indicate the modification ratio of each allele obtained from long-read sequencing, with gray gradient pairs representing data from two replicates.

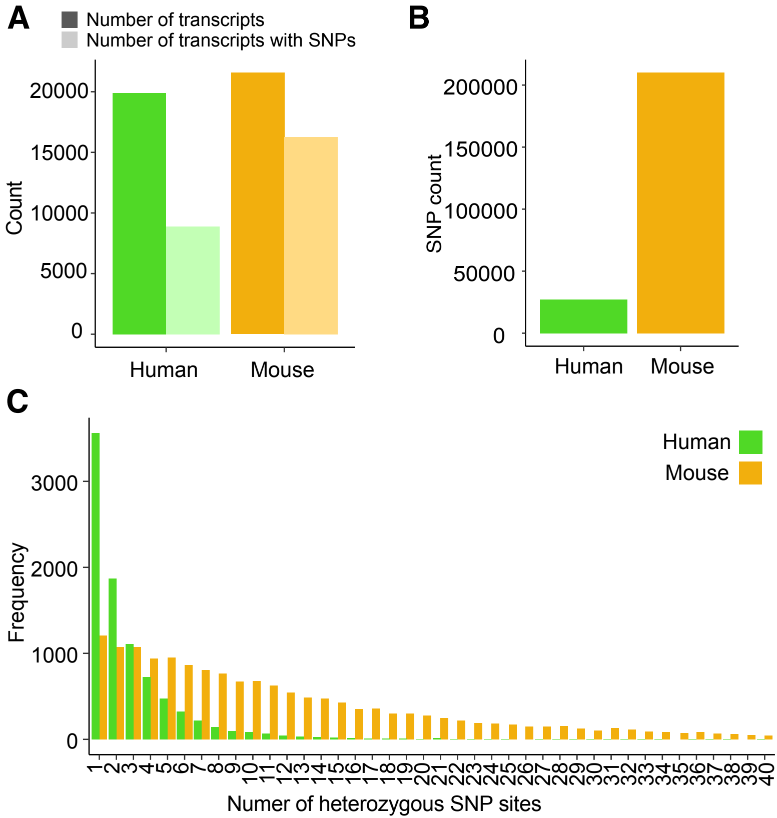


**Supplemental Figure. 8 | SNP counts in hybrid mESCs (C57BL/6J x CAST/EiJ) and NA12878.**

A) The number of transcripts between the two species was similar; however, transcripts with SNPs were significantly fewer in the NA12878 cell line. B) The number of heterozygous SNPs was significantly lower in NA12878, representing only one-tenth of the hybrid mESCs. C) Among the reads with at least one SNPs, 61% of NA12878 reads contained one or two SNPs per read, whereas 87% of the reads had more than two SNPs per read in hybrid mESCs.

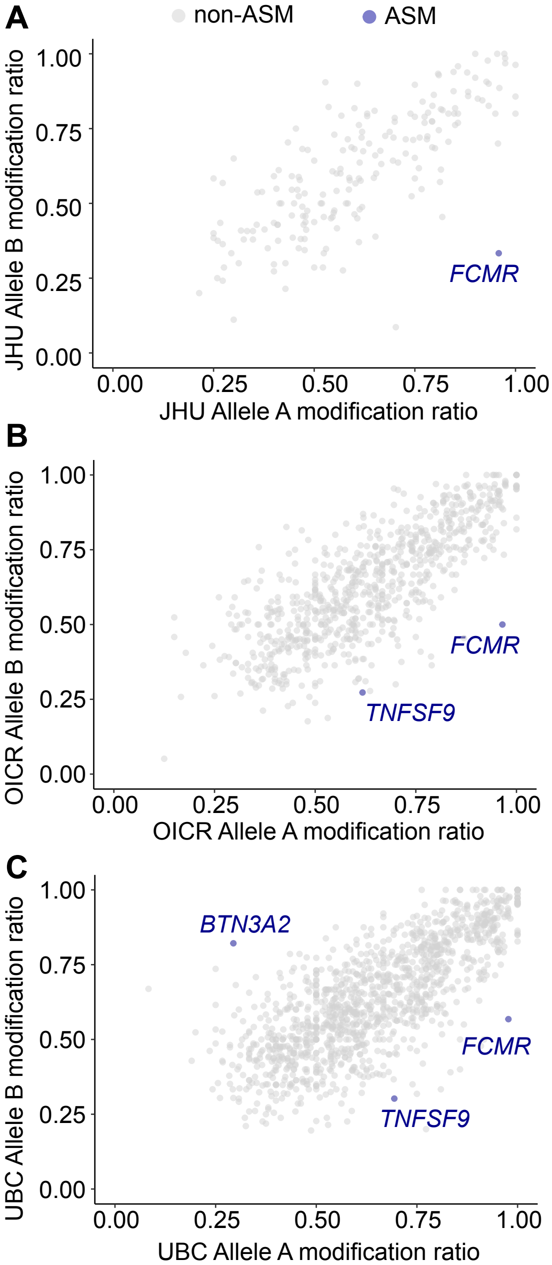


**Supplemental Figure. 9 | Allelic modification ratios in NA12878.**

Modification ratios from each allele, including three ASM sites (blue) and non-ASM sites (gray) in JHU (A), OICR (B) and UBC (C) datasets.

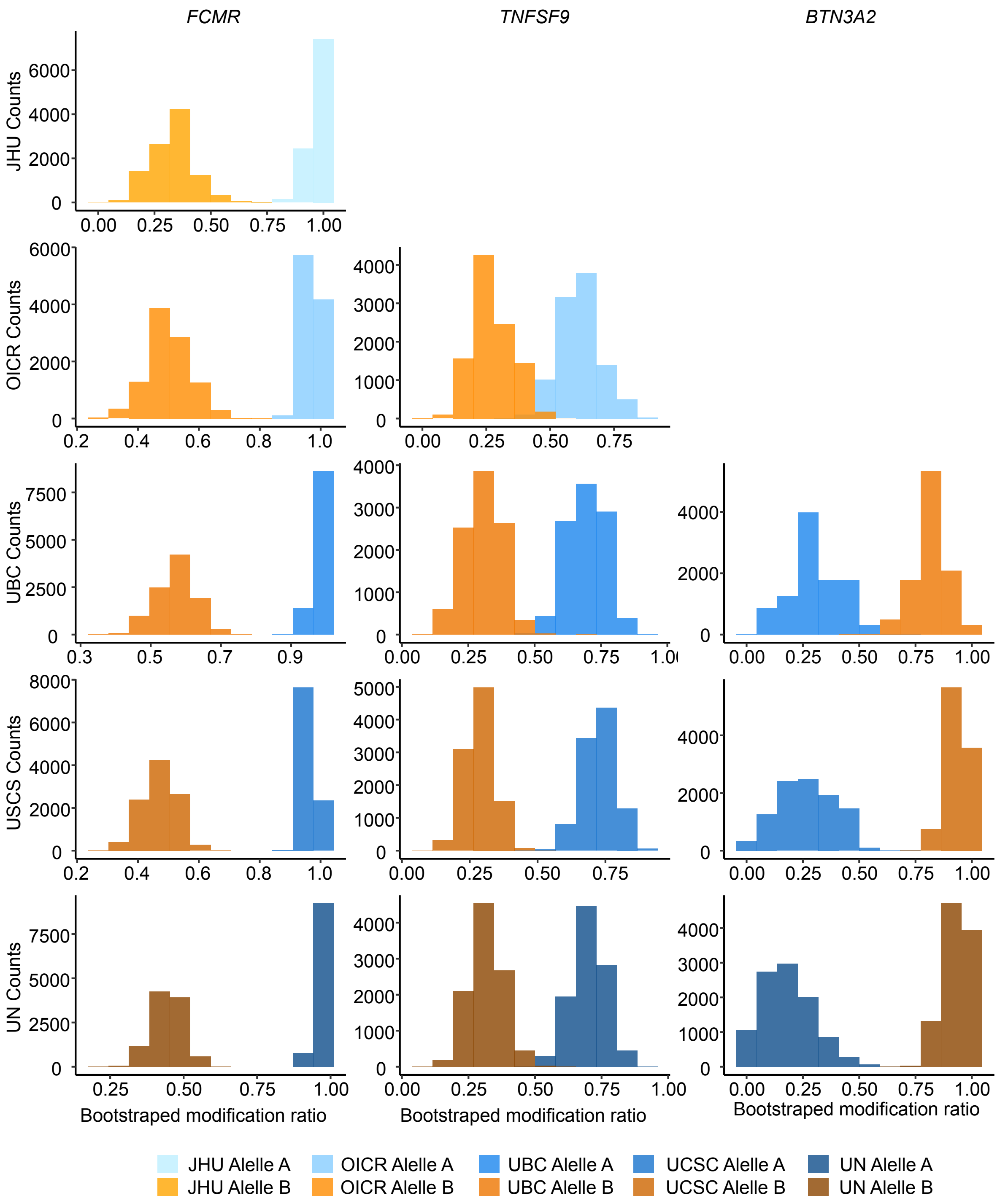


**Supplemental Figure. 10 | Bootstrapped modification ratios of ASM sites in NA12878.**

Resampled modification ratios from bootstrapping. Each color represents an allele (blue for Allele A, orange for Allele B), and the gray gradient indicates each replicate.

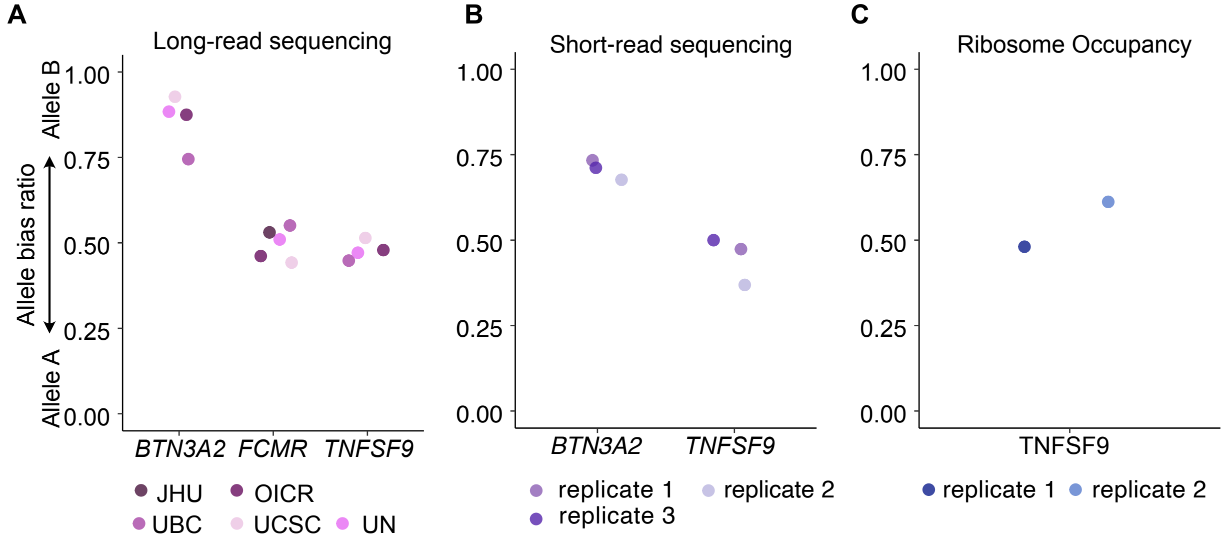


**Supplemental Figure. 11 | Allele specific RNA expression and ribosome occupancy of genes with ASM sites from NA12878**

Allele bias ratio of genes with ASM sites in NA12878. The allele bias ratio was calculated as [Allele B read counts / (Allele A read counts + Allele B read counts)]. Allele-specific RNA expression was measured from long-read (A) and short-read (B) sequencing. The allele-specific ribosome occupancy was detected by ribosome profiling (C).

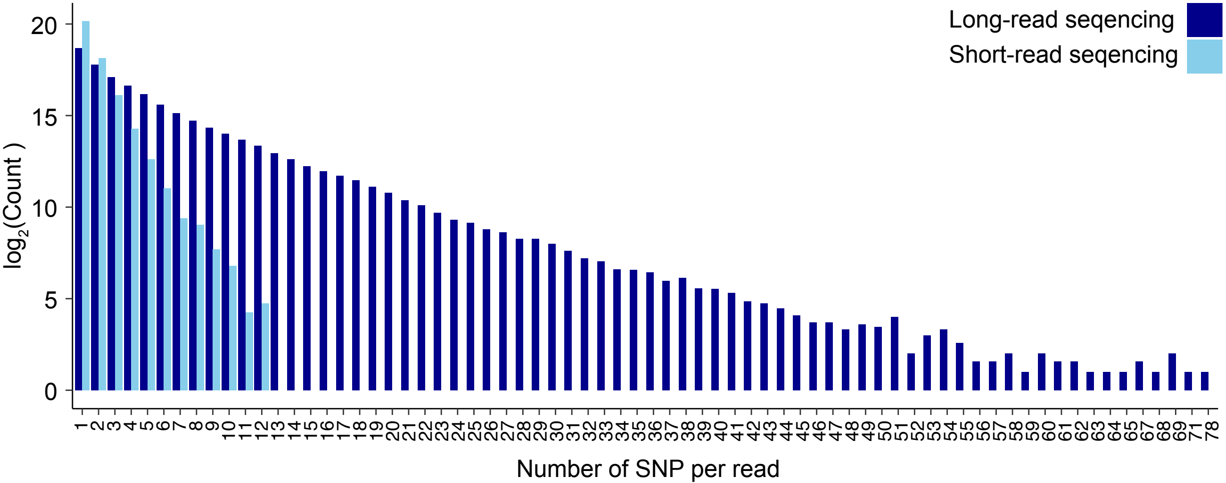


**Supplemental Figure. 12 | SNP counts per read**

SNP counts per read in short-read (lightblue) and long-read (dark blue) sequencing.

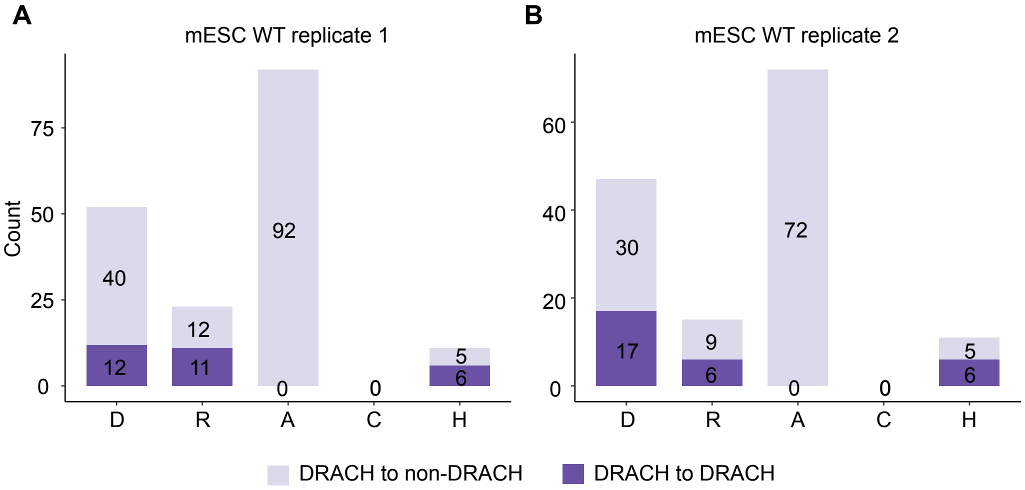


**Supplemental Figure. 13 | SNPs overlapping the DRACH motif**

SNPs can either convert DRACH motifs to non-DRACH motifs (light purple) or leave them matching another instance of the DRACH motif (dark purple) (A, replicate 1 and B, replicate 2).

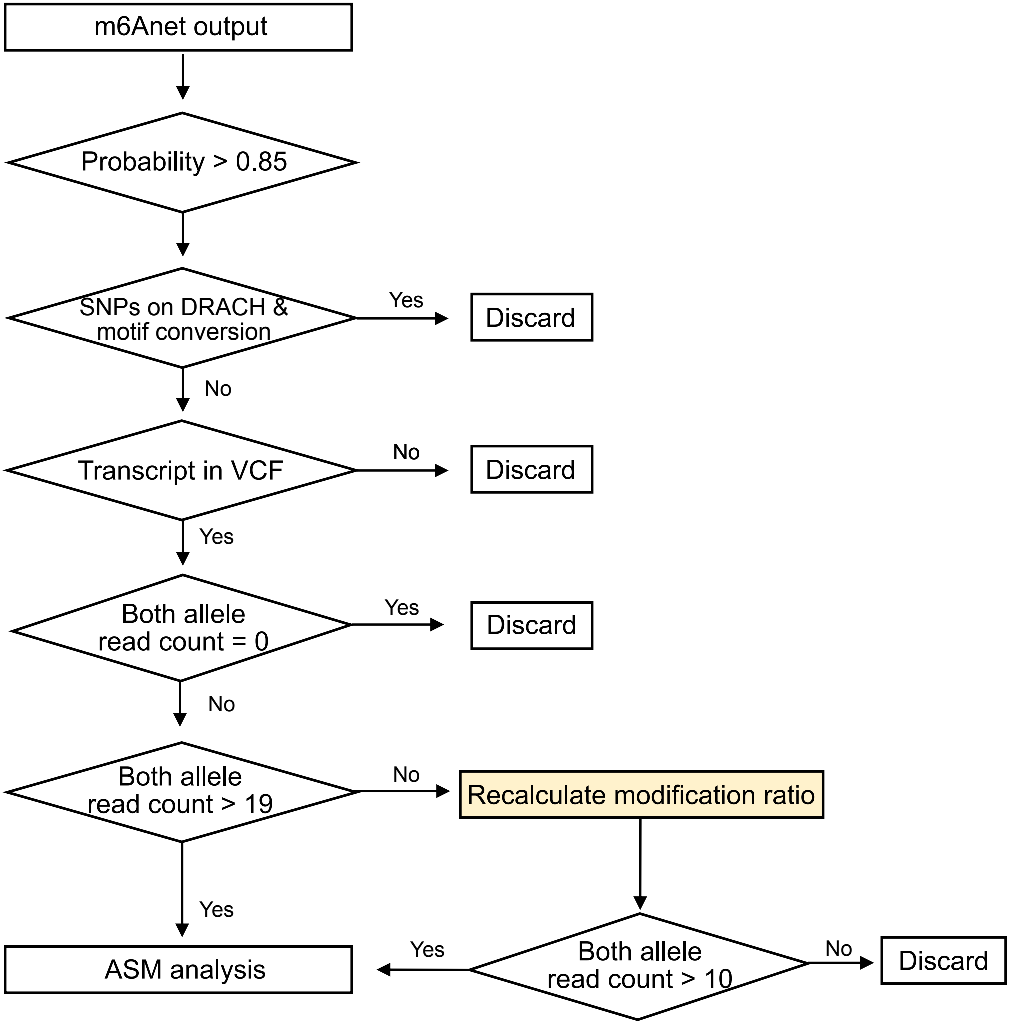


**Recalculation method:**

$$\text{Allele A modification ratio} = \frac{\text{(All read count * mod ratio from all reads) - (Allele B read count * Allele B mod ratio)}}{\text{Allele A read count}}$$

**Supplemental Figure. 14 | Data processing flowchart**
